# Repurposing Nelfinavir: AIM2 Inflammasome-Driven Anti-tumor Effects in Glioblastoma

**DOI:** 10.1101/2025.11.11.687819

**Authors:** Durgesh Meena, Shalini Chhipa, Priya Solanki, Lipika Sha, Sushmita Jha

**Author notes:** Corresponding author: Name: Sushmita Jha, Mailing address: Room no 303, Department of Bioscience and Bioengineering, Indian Institute of Technology Jodhpur, Jodhpur, Rajasthan 342037, India.

## Abstract

**Nelfinavir (NFR)**, originally developed as an antiretroviral agent for the human immunodeficiency virus, has demonstrated anti-cancer properties across various malignancies; however, its therapeutic potential in **Glioblastoma (GBM)** remains largely unexplored. In the present study, we investigated the anti-tumor effects of NFR in GBM using a comprehensive panel of experimental models, including established GBM cell lines, GBM cell line-derived spheroids, patient-derived primary glioma cells, and patient-derived glioma organoids. We further evaluated the synergistic potential of NFR in combination with standard chemotherapeutic agents, **Carboplatin** and **Doxorubicin,** across these platforms. *In vitro* analyses revealed that NFR significantly inhibits GBM cell proliferation and induces both apoptotic and necrotic cell death. Mechanistically, we identified activation of the **AIM2 inflammasome** as a potential additional pathway mediating the anti-tumor effects of NFR. Collectively, our findings highlight NFR as a promising therapeutic candidate for GBM, exerting its effects through anti-proliferative and pro-death mechanisms potentially linked to AIM2 inflammasome activation.

## Introduction

Gliomas represent the most common type of primary brain tumor in adults and are among the most aggressive and lethal forms of human cancer [1]. Gliomas originate from glial cells within the central nervous system (CNS). The World Health Organization (WHO) classifies adult-type diffuse gliomas into astrocytoma (grades 2, 3, or 4), oligodendroglioma (grades 2 or 3), and glioblastoma (grade 4) [1]. Glioblastoma (GBM) exhibits rapid growth, extreme cellular and molecular heterogeneity, diffuse infiltration into surrounding brain tissue, and a high rate of recurrence [2]. The current standard of care for newly diagnosed GBM follows a multimodal approach aimed at prolonging survival and preserving neurological function[3]. Initial management typically involves maximal safe surgical resection to reduce tumor burden and obtain tissue for diagnosis and molecular profiling. Postoperative treatment includes radiotherapy, most commonly delivered as 60 Gy combined with daily temozolomide (TMZ) chemotherapy [4,5]. Tumor Treating Fields (TTF), a non-invasive device-based therapy, may be considered in combination with TMZ for eligible patients, as it has shown statistically significant improvement in progression-free survival and overall survival [4,6]. Despite aggressive treatment involving surgical resection, radiotherapy, and chemotherapy, the median survival for GBM patients remains poor, typically ranging from 12 to 15 months following diagnosis. Therefore, there is a pressing need for novel therapeutic strategies to enhance survival outcomes in GBM patients.

Nelfinavir (NFR) is an FDA (Food and Drug Administration)-approved antiretroviral drug used to inhibit HIV (human immunodeficiency virus) replication. NFR treatment is associated with adverse metabolic effects in long-term treated patients, including hyperglycemia, insulin resistance, and lipodystrophy, suggesting mechanisms of action beyond its antiviral activity [7]. Notably, one proposed mechanism for NFR-induced insulin resistance involves the inhibition of the insulin-like growth factor 1 (IGF)/Akt (a serine/threonine kinase also called as PKB or protein kinase B) signaling pathway, which is frequently upregulated in various cancers [8]. This observation has sparked considerable interest in repurposing NFR as an anti-cancer agent. Drug repurposing strategies like this are particularly appealing because they can significantly shorten the drug development timeline and improve the affordability of cancer therapeutics for patients. NFR regulates multiple cellular pathways in mammalian cancer cells, but its primary molecular target responsible for its anti-tumor activity remains unclear[9]. Evidence from various studies suggests that the anti-cancer mechanisms of NFR may differ depending on the cancer type and cellular context. NFR induces cell death in glioma cell lines by activating the endoplasmic stress response, inhibiting proteasomal activity, and promoting the accumulation of misfolded proteins [10]. In breast cancer cells, reactive oxygen species (ROS) generation drives NFR-induced cytotoxicity [11]. In bladder (T24), head and neck (SQ20B), pancreatic (MIAPACA2), lung (A549), and rat fibroblast (REF) cancer cells, NFR inhibits Akt phosphorylation and enhances radiosensitivity both *in vitro* and *in vivo* [12]. NFR suppresses the proliferation of prostate cancer cells (LNCaP) by inhibiting androgen receptor (AR), STAT3 (signal transducer and activator of transcription 3), and AKT signaling pathways both *in vitro* and *in vivo* [13]. In melanoma cell lines, NFR inhibits growth by inducing apoptosis and G1 phase cell cycle arrest through CDK2 (cyclin-dependent kinase 2) inhibition and dephosphorylation of the retinoblastoma (Rb) tumor suppressor [14]. NFR also reduces angiogenesis *in vivo* in GBM cells through the downregulation of vascular endothelial growth factor (VEGF) and hypoxia-inducible factor 1-alpha (HIF-1α) [15]. Additionally, NFR induces growth arrest and apoptosis in non-small-cell lung cancer (NSCLC) cell lines (NCI-H460, NCI-H520, A549, EBC-1, and ABC-1), accompanied by suppression of Akt signaling [16]. In monocytes and macrophages (THP-1 and BMDMs), it has been shown that NFR disrupts the nuclear membrane by preventing the maturation of nuclear membrane proteins, including Lamin, leading to the release of nuclear DNA into the cytoplasm and subsequent activation of the AIM2 inflammasome [17]. These findings highlight the diverse mechanisms through which NFR affects various malignancies.

Several Phase I and II clinical trials have demonstrated the safety, tolerability, and potential therapeutic benefit of NFR in cancer patients, both as a monotherapy and in combination with other treatments, particularly in pancreatic cancer, NSCLC, and multiple myeloma (MM) [18–20]. However, most of these trials have been single-arm and open-label with small groups of patients. Therefore, larger randomized controlled trials are now needed to better evaluate the clinical effectiveness of NFR. In line with this, two large-scale randomized trials are currently underway to evaluate the effectiveness of NFR in combination with radiotherapy for treating locally advanced pancreatic cancer (NCT02024009) and cervical cancer (NCT03256916). A Phase I clinical trial investigating the combination of NFR, temozolomide, and radiotherapy in patients with GBM demonstrated that the treatment regimen is feasible and generally well tolerated [21]. However, the study was limited by a small cohort size and lacked a comprehensive molecular analysis of the tumors. Further investigations are required to delineate the anti-tumor properties of NFR and its underlying molecular mechanisms in GBM. In addition, well-designed Phase II clinical trials are needed to evaluate the safety and therapeutic efficacy of NFR in a larger, molecularly defined patient population [21].

In this study, we investigate the anti-tumor potential of NFR in GBM using a range of experimental models, including established GBM cell lines, spheroids, patient-derived primary glioma cells, and organoids. Given its reported anti-cancer activity in other malignancies, we examine NFR alone and in combination with standard chemotherapeutics, Carboplatin and Doxorubicin. Our data show that NFR inhibits proliferation and induces apoptotic and necrotic cell death in GBM cells. Importantly, we identify activation of the AIM2 inflammasome as a potential mechanism underlying NFR’s anti-tumor effects, supporting its promise as a therapeutic candidate for GBM.

## Material and Methods

### 1. Cell culture

The LN-18 (RRID: CVCL_0392) human glioblastoma cell line was obtained from the American Type Culture Collection (ATCC), while the LN-229 (RRID: CVCL_0393) and U87-MG (RRID: CVCL_0022) glioblastoma cell lines were sourced from the Cell Repository of the National Centre for Cell Science (NCCS), Pune, India. The human astrocyte cell line SVG was kindly provided by Dr Pankaj Seth from the National Brain Research Centre (NBRC), India. All cell lines were cultured in Dulbecco’s Modified Eagle Medium (DMEM; HiMedia, AL151A-500ML) supplemented with 10% fetal bovine serum (FBS; CellClone, CCS-500-SA-U) and 1% antibiotic-antimycotic solution containing penicillin, streptomycin, and amphotericin B (HiMedia, A002-200ML). Cultures were maintained at 37 °C in a humidified incubator with 5% CO₂.

### 2. Isolation and culture of primary cells from patient tissues

Glioma tissue samples were collected in artificial cerebrospinal fluid (aCSF) composed of 2 mM CaCl₂·2H₂O, 10 mM glucose, 3 mM KCl, 26 mM NaHCO₃, 2.5 mM NaH₂PO₄, 1 mM MgCl₂·6H₂O, and 202 mM sucrose at the All-India Institute of Medical Sciences (AIIMS), Jodhpur. Upon arrival at our laboratory, the aCSF was discarded, and the tissue was weighed. Samples were subsequently washed with fresh aCSF followed by PBS. The tissue was then minced into approximately 1 mm fragments and transferred into a 0.25% trypsin–EDTA solution. Mechanical dissociation was performed by shaking the samples at 250 RPM for 30 minutes at 37 °C. Following enzymatic digestion, a neutralizing medium was added to inactivate trypsin, and the suspension was centrifuged to collect the cells. The resulting cell pellet was resuspended in 1 ml of culture medium comprising 45% Dulbecco’s Modified Eagle Medium (DMEM), 45% F-12 nutrient mixture, and 10% fetal bovine serum (FBS). Cells were then seeded into tissue culture flasks and incubated in a humidified CO₂ incubator (5% CO₂, 37 °C, 95% humidity) [22].

### 3. Immunocytochemistry

Cells were seeded in chamber slides containing cell culture medium and incubated under standard conditions in a humidified CO₂ incubator (5% CO₂, 37 °C, 95% humidity). Following incubation, cells were washed with phosphate-buffered saline (PBS) and fixed with 4% paraformaldehyde (HiMedia, MB059-500ML) for 10 minutes at room temperature. Permeabilization was carried out using 0.1% Triton X-100 (Sigma, 1001723790) in PBS (permeabilization buffer) for 15 minutes. Subsequently, cells were blocked with blocking buffer consisting of 5% fetal bovine serum (FBS) in permeabilization buffer for 1 hour at 4 °C in a humidified chamber. For immunolabeling, cells were incubated overnight at 4 °C with the following primary antibodies: anti-AIM2 (rabbit; Sigma-Aldrich, Cat# SAB-4503648, RRID: AB_10752420), anti-GFAP (mouse; Cell Signaling Technology, Cat# 3670, RRID: AB_561049), anti-Lamin B1 (rabbit; Cell Signaling Technology, Cat# 13435, RRID: AB_2737428), and anti-HMGB1 (rabbit; Cell Signaling Technology, Cat# 3935, RRID: AB_2295241). The following day, slides were washed with PBS and incubated with the appropriate fluorescent secondary antibodies for 1 hour at 37 °C in a dark, humidified chamber: Alexa Fluor 488 (anti-rabbit; Invitrogen, Cat# A11008, RRID: AB_143165) and Alexa Fluor 594 (anti-mouse; Invitrogen, Cat# A11005, RRID: AB_2534073). Following incubation, slides were mounted for imaging using Fluoroshield^TM^ with DAPI (4’,6-diamidino-2-phenylindole) (Sigma, F6057-20ML)[23].

Microglia were identified by fixing cells with 4% paraformaldehyde for 10 minutes, followed by permeabilization using 0.1% Triton X-100 in PBS for 15 minutes at room temperature. Cells were then incubated with Ricinus communis agglutinin (RCA; Vector Laboratories, Cat# FL-1081, RRID: AB_2336708), a lectin marker for microglia, for 1 hour at 37 °C in a humidified chamber. Following PBS washes, cells were blocked in 5% FBS prepared in 0.1% Triton/PBS for 1 hour at 4 °C. Subsequently, cells were incubated overnight at 4 °C with a primary antibody against AIM2 (rabbit; Sigma-Aldrich, Cat# SAB-4503648, RRID: AB_10752420) in a humidified environment. The next day, cells were washed and incubated with Alexa Fluor 594-conjugated secondary antibody (anti-rabbit; Life Technologies, Cat# A11012, RRID: AB_2534079) for 1 hour at 37 °C in a dark, humidified chamber. After final washes with PBS, cells were mounted as previously described [24].

### 4. Western blotting

Cells or tissues were lysed in radioimmunoprecipitation assay (RIPA) buffer supplemented with freshly added protease inhibitors and incubated for 4 minutes at 4 °C. Protein concentrations were quantified using the Bradford assay. Equal amounts of protein (15 µg per sample) were resolved by SDS-PAGE and subsequently transferred onto nitrocellulose membranes. Membranes were blocked with 5% skimmed milk in TBST for 1.5 hours at room temperature, followed by overnight incubation at 4 °C with the following primary antibodies: anti-AIM2 (rabbit; Sigma-Aldrich, Cat# SAB-4503648, RRID: AB_10752420), anti-ASC (rabbit; Santa Cruz Biotechnology, Cat# sc-22514-R, RRID: AB_2174874), anti-Caspase-1 (mouse; Santa Cruz Biotechnology, Cat# sc-56036, RRID: AB_781816), anti-Lamin B1 (rabbit; Cell Signaling Technology, Cat# 13435, RRID: AB_2737428), anti-HMGB1 (rabbit; Cell Signaling Technology, Cat# 3935, RRID: AB_2295241), and anti-β-actin (mouse; Santa Cruz Biotechnology, Cat# sc-47778, RRID: AB_626632). The membranes were then incubated with appropriate HRP-conjugated secondary antibodies (anti-rabbit IgG, Cell Signaling Technology, Cat# 7074, RRID: AB_2099233; anti-mouse IgG, Cell Signaling Technology, Cat# 7076, RRID: AB_330924). Protein bands were visualized using the Azure Biosystems gel documentation system. Densitometric analysis was performed using ImageJ (RRID: SCR_003070) software, and relative protein levels were normalized to β-actin expression followed by untreated control [25].

### 5. Colony formation assay

UT and NFR (4μM) treated GBM (U87-MG) cells were seeded at a density of 2500 cells per well (5% CO₂; 37°C) in a chamber slide, and small colonies were observed after 36–48 hours [23]. Slides were observed using a bright field microscope, and results were quantified by counting the number of colonies formed per well and cells present per colony.

### 6. Spheroid formation

GBM (U87-MG) and astrocyte (SVG) cells were seeded in low-attachment U-shaped 96-well plate for spheroid generation. Cells were maintained in a medium containing 0.1% antibiotic,10% FBS, and nutrient media (DMEM) at 37°C with 5% CO₂. Cells began to aggregate within a few hours and formed spheroids within one day. The spheroids were imaged with a multimode reader (cytation5, Agilent) [26].

### 7. Image acquisition of spheroid

Imaging of seeded spheroids was performed using an automated multimode reader (Cytation 5, Agilent) starting from day zero under controlled conditions (5% CO₂, 37 °C). Images were captured at multiple Z-planes to ensure comprehensive visualization of the three-dimensional structures. Following acquisition, Z-stack images were processed and stitched using the instrument’s integrated software. The final composite images were saved for subsequent analysis [26].

### 8. Quantification of area and diameter of spheroids

Spheroid diameter and area were quantified using NIH ImageJ software. The spheroid area was manually outlined and measured using the application’s built-in tools. Diameter measurements were taken along two perpendicular axes—horizontal and vertical—and the average of these two values was reported as the spheroid diameter [27] [26].

### 9. Quantification of circularity and compactness of spheroids

Blinded evaluators assessed spheroid circularity and compactness using a 5-point scoring system. For both categories, the absence of a discernible spheroid or the presence of multiple spheroids was assigned a score of 1. Spheroids displaying visible spaces and gaps were classified as loose aggregates (score = 2), while those lacking internal gaps but exhibiting diffuse borders were rated as tight aggregates (score = 3). Further compaction led to the formation of pronounced dark borders around spheroids with minimal loosely connected cells, categorized as compact spheroids (score = 4)[26]. The highest degree of compaction was characterized by surface cells closely conforming to the spheroid contour, producing a smooth and well-defined outline, designated as tight spheroids (score = 5). For circularity, spheroids with a balanced presence of concave and convex edges were classified as irregular (score = 2). Those predominantly convex with minor concave indentations were rated as minor irregular (score = 3). Elongated spheroids lacking concave sections received a score of 4, while perfectly symmetrical, circular spheroids were assigned a score of 5 [27].

### 10. Phalloidin Staining

GBM and astrocyte spheroids were treated with NFR (7μM, 14μM) for 48h hours. After the treatment, spheroids were washed with PBS, fixed using 4% paraformaldehyde (HiMedia, MB059-500ML) for 10 minutes, and permeabilized with 0.1% triton-x (Sigma, 1001723790) in PBS for 15 minutes. Spheroids were stained with Rhodamine phalloidin (Invitrogen, R415) solution in 0.1% triton-x (1:200) for 2 hours at RT. Spheroids were washed with PBS and mounted using Fluoroshield^TM^ with DAPI (4’,6-diamidino-2-phenylindole) (Sigma, F6057-20ML) [28].

### 11. Flow Cytometry analysis of Annexin-PI-stained spheroids

U87-MG spheroids were treated with NFR (7μM, 14μM) for 48 hours. After the treatment, as per the manufacturer’s protocol (Invitrogen™, Cat#V13242), spheroids were washed with PBS, trypsinized for 10 minutes, and then spheroids were resuspended in 100μl of Annexin V Binding Buffer. Each sample was incubated with 5 μl of fluorescein isothiocyanate (FITC)-conjugated Annexin V and 1μl of 100 μg/ml PI for 15 min at room temperature in the dark. Finally, 400 μl of Annexin V Binding Buffer was added to each sample and acquired with Guava^®^ easyCyte^™^ flow cytometer [11].

### 12. Fluorescence microscopy of Annexin-PI-stained spheroids

U87-MG spheroids were treated with NFR (7μM, 14μM) for 48 hours. After the treatment, as per the manufacturer’s protocol (Invitrogen™, Cat#V13242), spheroids were washed with PBS and incubated with 5 μl of fluorescein isothiocyanate (FITC)-conjugated Annexin V and 2μl of 100 μg/ml PI in 100μl of Annexin V Binding Buffer for 15 min at room temperature in the dark. Subsequently, spheroids were washed with Annexin V Binding Buffer, placed on a glass slide, and mounted.

### 13. MTT assay for spheroids

Spheroids were treated with NFR (4μM), Carboplatin (2.7mM), or Doxorubicin (4.9μM) for 48 hours. After the treatment, spheroids were collected in an Eppendorf tube, washed with PBS, trypsinized for 15 minutes, and then resuspended in 90μl DMEM. These DMEM resuspended cells were transferred back to a 96-well plate, and 10μl of MTT solution was added to each well. The plate was kept in an incubator (Model 170S, Eppendorf) at 37°C with 5% CO₂ in the dark for 2 hours. After incubation, 100μl of acidic isopropanol solution was added to each well and mixed thoroughly using a pipette. Absorbance at 570nm was measured using a multi-mode microplate reader (Synergy H1 Hybrid, Biotek Instruments Inc). The IC_50_ value of the NFR was calculated from the cell viability data [29].

### 14. MTT assay for 2D culture

10,000 cells were seeded per well in a 96-well plate and were incubated at 37°C with 5% CO₂. Cells were treated with different concentrations of NFR, Carboplatin, or Doxorubicin. After the treatment, the drug-containing media was removed carefully from the microplate. 100μl of fresh serum-free media and 10μl of MTT solution (Sigma) were added to each well. The plate was kept in an incubator (Model 170S, Eppendorf) at 37°C with 5% CO₂ in the dark for 2 hours. After incubation, 100μl of acidic isopropanol solution was added to each well and mixed thoroughly using a pipette. Absorbance at 570nm was measured using a multi-mode microplate reader (Synergy H1 Hybrid, Biotek Instruments Inc). IC_50_ value of the NFR was calculated from the cell viability data [30].

### 15. Statistical Analyses

Data are presented as mean ± SEM. Unmatched Student’s t-tests were employed to statistically assess significant differences. Differences were deemed statistically significant if p<0.05. The student’s t-test was conducted to ascertain significant differences between groups.

## Results

### 1. Nelfinavir selectively reduces the viability and proliferation of Glioblastoma cells

To determine the effect of NFR on the viability of GBM cells, we performed an *in vitro* assay on U87-MG cells, treated with NFR. We also took normal human astrocyte cells as controls. The cells were treated with 5, 10, 15, 20, 25, and 30μM NFR for 24 hours. This concentration is attainable in the plasma of human patients who take NFR at a standard dosage [31]. As shown in Figure 1A, B, NFR decreases the viability of GBM and astrocyte cells, in a dose-dependent manner. The IC_50_ concentration of NFR in GBM cells is 5.15μM, and for astrocytes, 4.2μM. Further, we performed a colony formation assay to test the anti-proliferative properties of NFR in GBM cells. As shown in Figure 1C-D, NFR-treated GBM cells show a significant decrease in the number of colonies. This data confirms the anti-proliferative properties of NFR in GBM cells. This aligns with already published data in melanoma, breast cancer, and glioma cells [32–34].

**Legend Figure 1.**
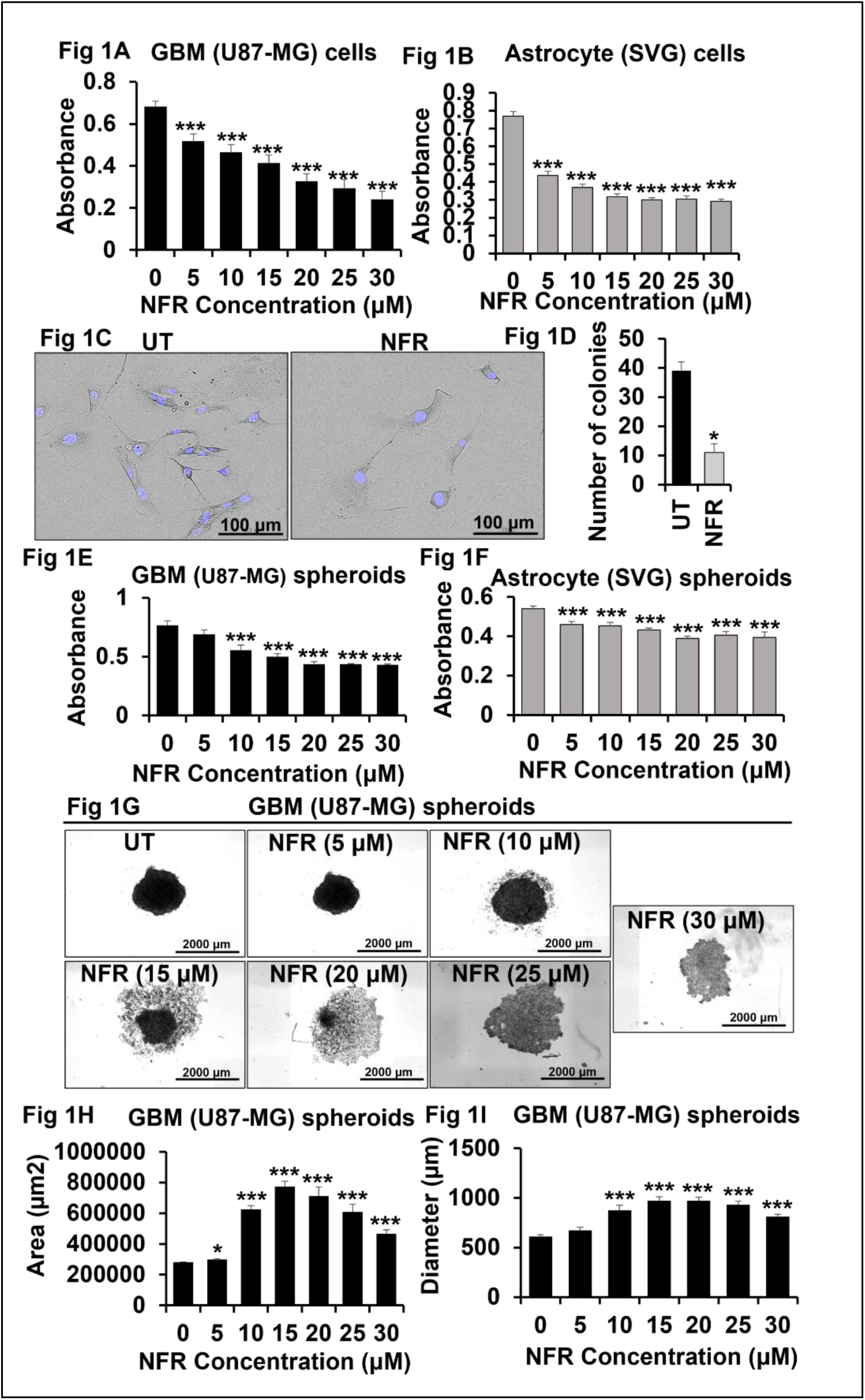
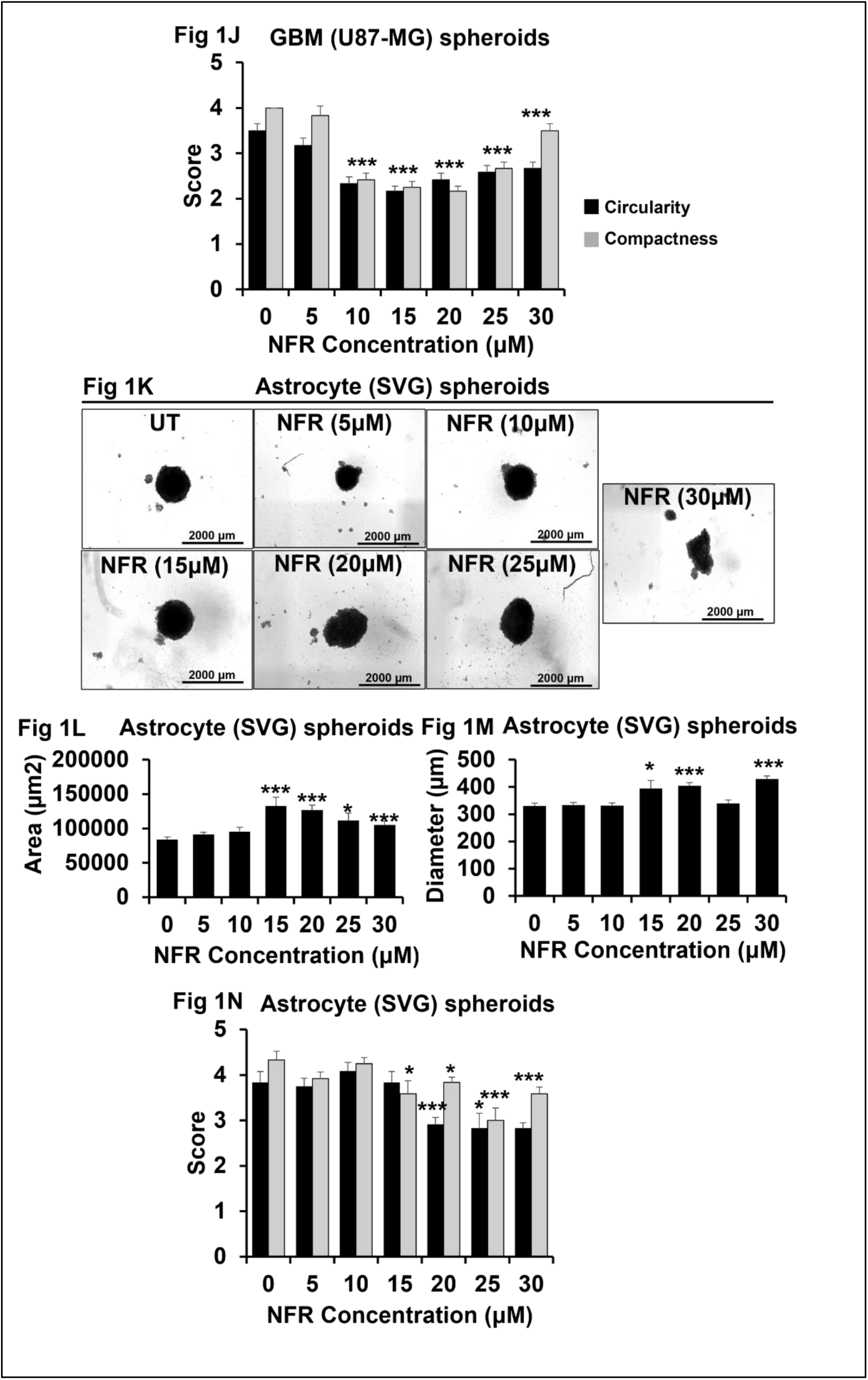
Nelfinavir reduces the viability and proliferation of Glioblastoma cells: A, B. Cell viability assay was performed in GBM (U87-MG) and astrocyte (SVG) cells treated with NFR at indicated concentrations for 24 hours. The graph is representative of three experiments. C-F. Colony formation assay was performed in UT and NFR (4μM, 48 hours) treated U87-MG cells. C. DAPI was used to stain nuclei (blue). At least 50 frames were imaged per well of cells cultured in two-well chamber slides, Scale bar, 100μm. Images are representative of 3 experiments. D. The number of colonies was counted in UT and NFR-treated U87-MG cells by a blind reader. The experiment was performed three times. The graph presents data from one representative experiment. E, F. Cell viability assay was performed in GBM (U87-MG) and astrocyte (SVG) spheroids treated with NFR at indicated concentrations for 48 hours. The graph is representative of 2 experiments. G, K. Brightfield images of GBM (U87-MG) and astrocyte (SVG) spheroids treated with NFR. Images are representative of two experiments. Scale bar, 2000μm. H, I, L, M. The area and diameter of GBM (U87-MG) and astrocyte (SVG) spheroids were quantified using ImageJ. The graph is representative of two experiments. J, N. Circularity and compactness of U87-MG and SVG spheroids were analysed by blind readers. The graph is representative of two experiments. Data are represented as mean ± SEM. Error bars indicate SEM. ^∗^p < 0.05, **p < 0.01, ***p < 0.005 (Student’s t-test).

Building on our findings that NFR regulates the viability and proliferation of GBM cells in 2D (two-dimensional) cultures, we next examined its effect on 3D (three-dimensional) tumor growth. 3D cultures provide a more physiologically relevant model compared to traditional 2D systems. Unlike 2D cultures, 3D tumor spheroids better mimic the *in vivo* GBM microenvironment, particularly in terms of spatial cell-cell and cell-extracellular matrix (ECM) interactions [35] [36]. These spheroids arise through spontaneous cell aggregation. Following initial cell-cell contact, GBM cells upregulate E-cadherin, which accumulates on the cell surface. Strong intercellular E-cadherin interactions then drive the formation of compact spheroidal structures [37]. To further explore the effect of NFR in a more physiologically relevant context, we generated spheroids using both GBM (U87-MG) and astrocyte (SVG) cells. Once the spheroids were fully formed, they were treated with 4μM NFR (Figure S1A, B). This concentration was selected based on the IC_50_ values of NFR determined for both GBM and astrocyte cells in 2D culture. At this concentration, there was no noticeable difference in GBM and astrocyte cell viability between untreated and NFR-treated spheroids (Figure S1A, B). This prompted the question of whether NFR is inherently ineffective in 3D culture conditions, or if the differences between 2D and 3D microenvironments necessitate a different concentration for efficacy. To address this, GBM and astrocyte spheroids were treated with a range of NFR concentrations, including 5, 10, 15, 20, 25, and 30μM NFR. As shown in Figure 1E-F, NFR decreases the viability of GBM and astrocyte spheroids in a dose-dependent manner. Notably, the effective dose required to reduce viability in 3D spheroids was significantly higher compared to 2D cultures. Moreover, the IC_50_ concentration for GBM spheroids was significantly lower than that for astrocyte spheroids, indicating that GBM cells are more sensitive to NFR in a 3D microenvironment. These findings contrast with our earlier observations in 2D cultures, where the IC_50_ concentrations of NFR for GBM and astrocyte cells were nearly equivalent. Microscopic imaging further supported these findings (Figure 1G, K) (Figure S1C). NFR treatment led to the disintegration of GBM spheroids, with cells visibly detaching and migrating out of the spheroid structure. In contrast, astrocyte spheroids maintained their structural integrity with minimal disruption. To quantify these observations, the area and diameter of spheroids were measured using ImageJ. Figures 1H and 1I show a significant increase in the area of GBM spheroids starting at 5μM NFR, with a corresponding increase in diameter observed at 10μM. In comparison, astrocyte spheroids exhibit minimal changes at these concentrations (Figure 1L, 1M). Conversely, astrocyte spheroids display a significant increase in area and diameter only following treatment with 15μM and 20μM NFR, respectively (Figure 1L, M). Notably, the observed increase in spheroid size does not indicate cellular growth, as our MTT data clearly demonstrate substantial cell death at these concentrations [38]. Rather, the enlarged area reflects spheroid disintegration, with dead or dying cells dispersing from the core, thereby contributing to the apparent expansion in size. These findings further indicate that NFR induces cell death in GBM spheroids at significantly lower concentrations compared to astrocyte spheroids. Additionally, actin and DNA staining were performed to visualize the cytoskeleton and nuclei of NFR-treated GBM and astrocyte spheroids (Figure S1D, E). The fluorescent images revealed significant structural disruption and large gaps within NFR-treated GBM spheroids, whereas astrocyte spheroids remained compact and intact (Figure S1A, B). Furthermore, we assessed the circularity and compactness of NFR-treated GBM and astrocyte spheroids to get insight into the impact of NFR on 3D structure of the tumor (Figure 1J and 1N). Figures 1J and 1N show that GBM spheroids exhibit a significant reduction in circularity and compactness at lower NFR concentrations compared to astrocyte spheroids, indicating greater structural disruption in the GBM spheroids. These disruptions further indicate the loss of cellular integrity and death of cells in GBM spheroids. Previously, in prostate cancer spheroids, it has been reported that as the treatment progresses, cell-to-cell and cell-to-matrix interactions are disrupted due to cytotoxicity, thus leading to disruption of cell aggregation [39]. In summary, these findings indicate that NFR exhibits greater selectivity and efficacy in targeting GBM cells over normal cells in a 3D environment.

### 2. Synergistic Anti-tumor Effects of Nelfinavir with Carboplatin and Doxorubicin

GBM is a highly aggressive tumor and is currently managed using a multimodal treatment strategy that includes surgical resection, radiotherapy, and chemotherapy with temozolomide [5]. Alternatives to current therapies, recent studies show that Carboplatin and Doxorubicin could have a significant potential for improved GBM treatments [40,41]. Doxorubicin exerts its anti-cancer effects through multiple mechanisms [42]. It intercalates into DNA, blocking replication and transcription, and inhibits topoisomerase II, leading to DNA double-strand breaks and apoptosis. It also generates reactive oxygen species (ROS), causing oxidative damage to DNA, proteins, and lipids. On the other hand, Carboplatin acts as an alkylating agent, causing cross-linking between and within DNA strands, leading to inhibition of DNA, RNA, and protein synthesis and triggering programmed cell death, mostly in rapidly dividing cells [43]. We evaluated the effect of NFR in combination with the chemotherapeutic agents Carboplatin and Doxorubicin on GBM and astrocyte cells. In this direction, cells were treated individually with NFR (4μM), Carboplatin (2.7mM), or Doxorubicin (4.9μM), as well as with combinations of NFR and Carboplatin and NFR and Doxorubicin. Figure 2A and B show that the individual effects of NFR were comparable to those of Carboplatin and Doxorubicin in 2D culture. However, when combined, NFR significantly enhanced the cytotoxicity of both chemotherapeutic agents, leading to a more pronounced reduction in cell viability compared to treatment with NFR, Carboplatin, or Doxorubicin alone. These findings suggest a synergistic interaction between NFR and the chemotherapeutics (Carboplatin, Doxorubicin), enhancing their overall efficacy. Representative images in Figure 2C further illustrate the distinct patterns of cell death induced by these treatments. While Carboplatin and Doxorubicin predominantly induced apoptosis, NFR treatment led to a combination of apoptotic and necrotic cell death. Similarly, in the combinatorial treatments, cells exhibited features of both apoptosis and necrosis, indicating a broader cytotoxic effect (Figure 2C). Earlier studies have also shown that Carboplatin and Doxorubicin induce apoptosis in GBM cells [40,41].

**Legend Figure 2.**
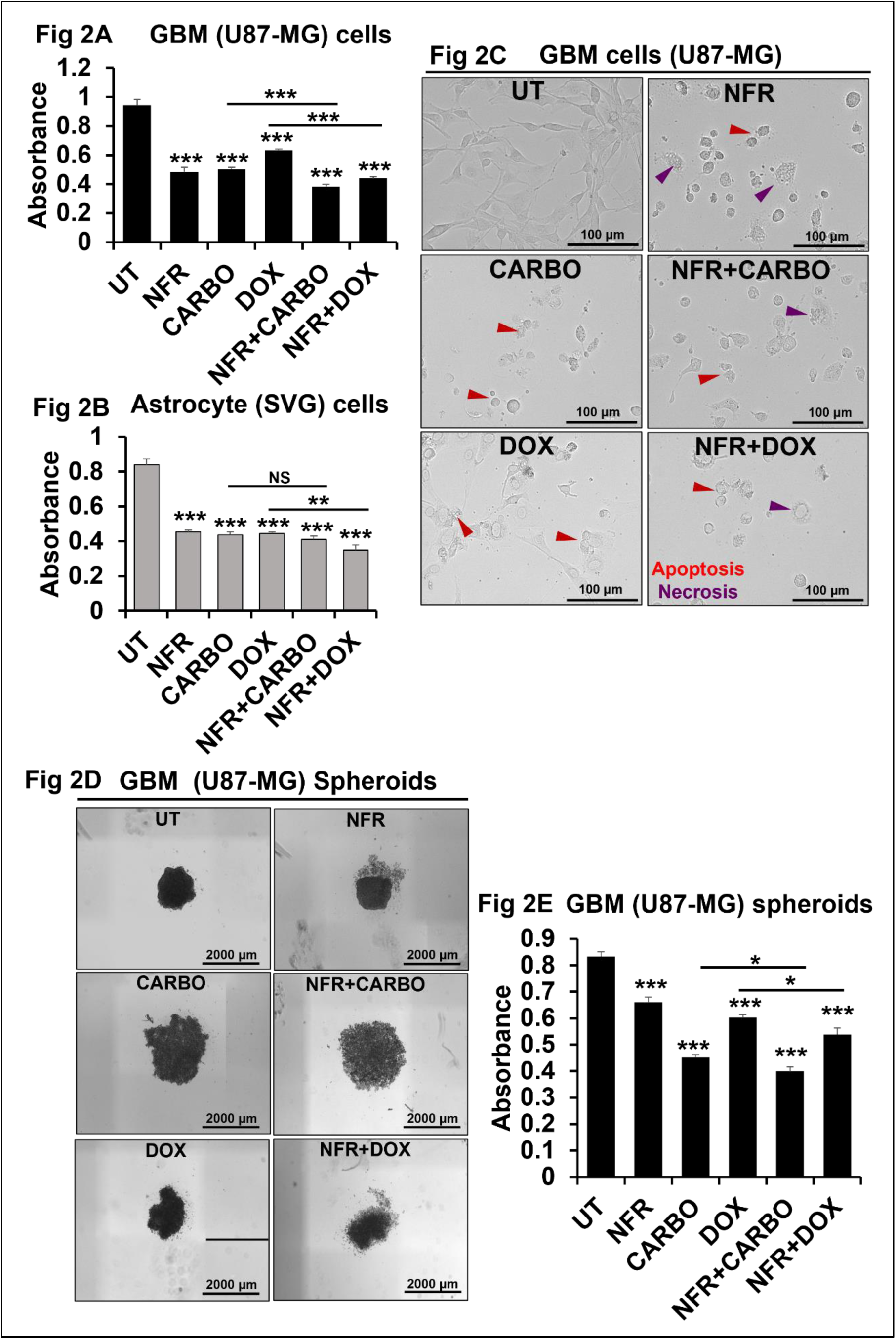
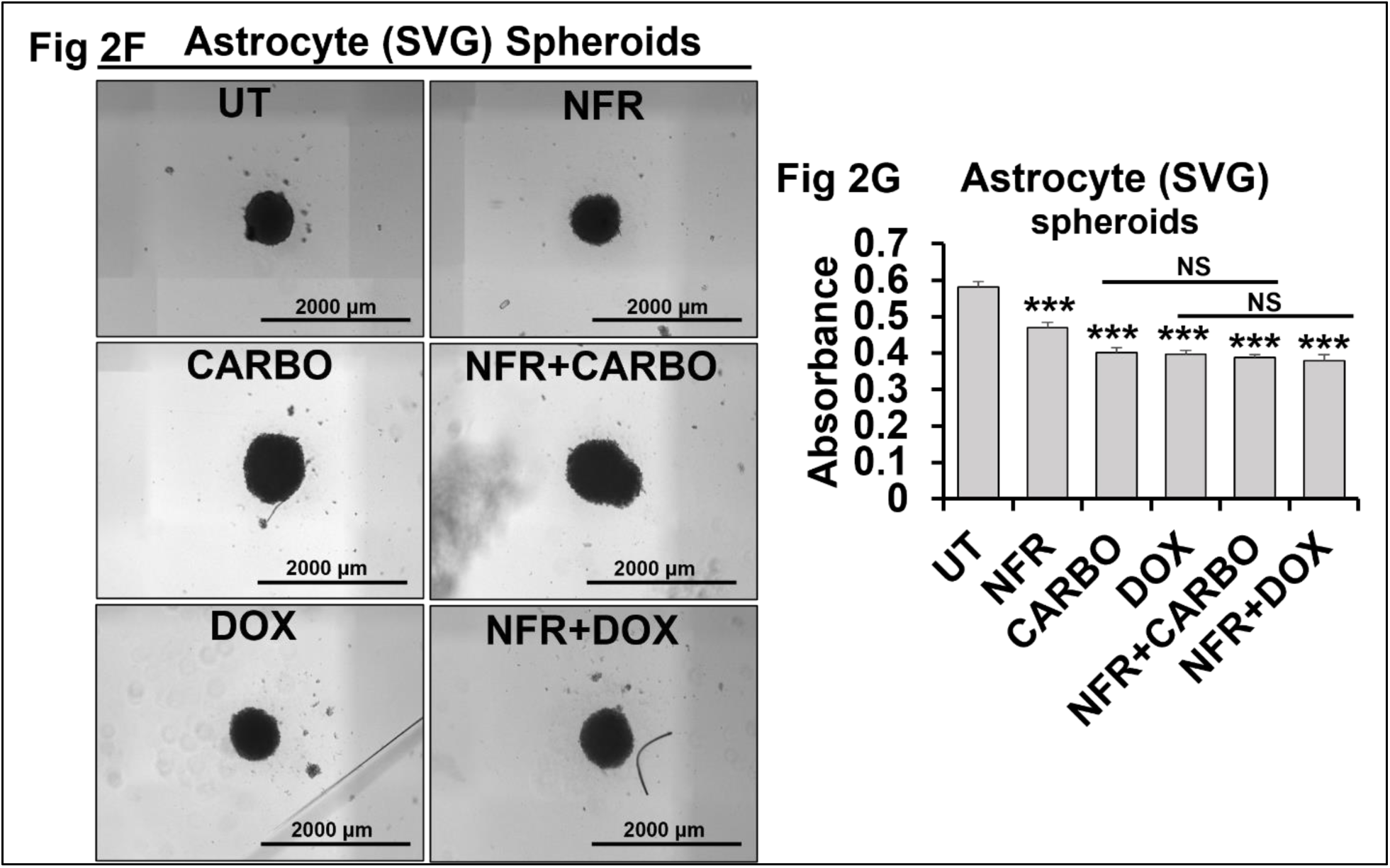
Synergistic Anti-Tumor Effects of Nelfinavir with Carboplatin and Doxorubicin: A, B. Cell viability assay was performed in GBM (U87-MG) and astrocyte (SVG) cells treated with NFR (4μM), Carboplatin (CARBO, 2.7mM), Doxorubicin (DOX, 4.9μM), or in indicated combination for 48 hours. The graph is representative of three experiments. C. Brightfield images of GBM (U87-MG) and astrocyte (SVG) cells treated with NFR, CARBO, DOX, or in indicated combination for 48 hours. Images are representative of 3 experiments. Scale bar, 100μm. E, G. Cell viability assay was performed in GBM (U87-MG) and astrocyte (SVG) spheroids treated with NFR (7μM), CARBO (2.7mM), DOX (4.9μM), or in indicated combination for 48 hours. The graph is representative of three experiments. D, F. Brightfield images of GBM (U87-MG) and astrocyte (SVG) spheroids treated with NFR, CARBO, DOX, or in indicated combination for 48 hours. Images are representative of 2 experiments. Scale bar, 2000μm. Data are represented as mean ± SEM. Error bars indicate SEM. * p < 0.05, **p < 0.01, ***p < 0.005 (Student’s t-test).

Further to extend these observations in the 3D microenvironment, we examined the impact of NFR and its combinations with Carboplatin and Doxorubicin in 3D spheroid cultures derived from GBM and astrocyte cells. From figures 2E, 2G, it is evident that NFR (7μM) alone significantly reduced cell viability in both GBM and astrocyte spheroids. However, when combined with chemotherapeutic agents, Carboplatin (2.7 mM) and Doxorubicin (4.9μM), NFR markedly enhanced their efficacy in GBM spheroids, while having minimal additional impact on astrocyte spheroids (Figure 2E, G) (Figure S2C). These findings highlight that in the 3D microenvironment, NFR potentiates the effects of conventional chemotherapies, Carboplatin and Doxorubicin, specifically in malignant GBM cells. Figures 2D and 2F present representative images showing that the combination of NFR with Carboplatin or Doxorubicin results in enhanced toxicity toward GBM spheroids, while having minimal impact on normal astrocyte spheroids. These results further support the conclusion that NFR acts synergistically with Carboplatin and Doxorubicin to target and eliminate GBM cells selectively. On the other hand, treatment with NFR (4μM) in combination with either Carboplatin or Doxorubicin did not enhance the efficacy of these drugs (Figure S2A-B).

### 3. Nelfinavir induces Apoptosis and Necrosis in Glioblastoma cells

The type of cell death induced by NFR is cell type and context-dependent. In melanoma (WM115), prostate cancer (LNCaP), NSCLC (NCI-H460), and hepatocellular carcinoma (HepG2 and WCH-17) cell lines, NFR predominantly triggers apoptosis [34,44–46]. In contrast, in breast cancer cell lines such as MDA-MB-231 and MCF, NFR induces both apoptosis and necrosis [32]. Additionally, in GBM cells (U251), one of the reported mechanisms by which NFR induces cell death involves activation of the endoplasmic reticulum stress response (ESR) [33]. Based on previous studies, we investigated whether NFR induces apoptosis or necrosis in GBM cells. To this end, we performed Annexin V-FITC and propidium iodide (PI) staining on NFR-treated GBM spheroids. Annexin V is a protein that binds specifically to phosphatidylserine, a membrane phospholipid typically localized to the inner leaflet of the plasma membrane but externalized during early apoptosis. PI is a membrane-impermeant DNA-binding dye that can only enter cells with compromised membranes, thus staining late apoptotic or necrotic cells [47]. Based on their staining profiles, cells can be classified as: healthy (Annexin V^−^ / PI^−^), early apoptotic (Annexin V^+^ / PI^−^), late apoptotic (Annexin V^+^ / PI^+^), or necrotic (Annexin V^−^ / PI^+^). As shown in Figure 3A, at a lower NFR concentration (7μM), GBM cells predominantly exhibit Annexin V staining (green fluorescence), indicating early apoptosis, with only a few cells showing PI staining (red fluorescence), suggestive of necrosis. In contrast, treatment with a higher concentration of NFR (14μM) results in a marked increase in PI fluorescence (red) intensity and a greater number of PI-positive cells (Figure 3A) (Figure S3A-D). These observations indicate that NFR induces both apoptotic and necrotic cell death in GBM cells in a concentration-dependent manner. Since PI staining can also indicate late-stage apoptosis, we further assessed the type of cell death by examining HMGB1 (High Mobility Group Box 1) expression. HMGB1 is a nuclear protein that normally remains tightly bound to chromatin [48]. However, during necrosis, the loss of nuclear membrane integrity leads to the release of HMGB1 into the cytoplasm. This translocation is a hallmark of necrotic cell death and is often accompanied by inflammation, as extracellular HMGB1 functions as a damage-associated molecular pattern (DAMP). Therefore, elevated HMGB1 levels, particularly with cytoplasmic localization, are widely used as an indicator of necrosis, distinguishing it from apoptosis, where HMGB1 remains localized within the nucleus. To investigate this, GBM cells were treated with increasing concentrations of NFR (4μM, 10μM, and 20μM), followed by western blot analysis for HMGB1 (Figure 3B, 3C). The results showed a dose-dependent increase in HMGB1 expression with higher NFR concentrations. This upregulation of HMGB1 further supports the induction of necrotic cell death in response to NFR treatment in GBM cells. Furthermore, we conducted fluorescence microscopy to assess the subcellular localization of HMGB1 in NFR-treated GBM cells (Figure 3D) (Figure S3D). The fluorescent images reveal the cytoplasmic localization of HMGB1 in NFR-treated non-viable GBM cells, further indicating necrotic cell death. To further quantify the proportion of apoptotic and necrotic cells, we performed flow cytometry on Annexin V-FITC and PI-stained GBM spheroids treated with NFR (7μM, 14μM) (Figure 3E, F). The flow cytometry data clearly show that upon increasing NFR concentrations, the number of apoptotic and necrotic cells increases in GBM spheroids (Figure 3E, 3F). These results further confirm that NFR induces GBM cell death in a dose-dependent manner. These findings are consistent with previous observations in breast cancer cells, where NFR treatment has been shown to induce both apoptotic and necrotic cell death [32].

**Legend Figure 3.**
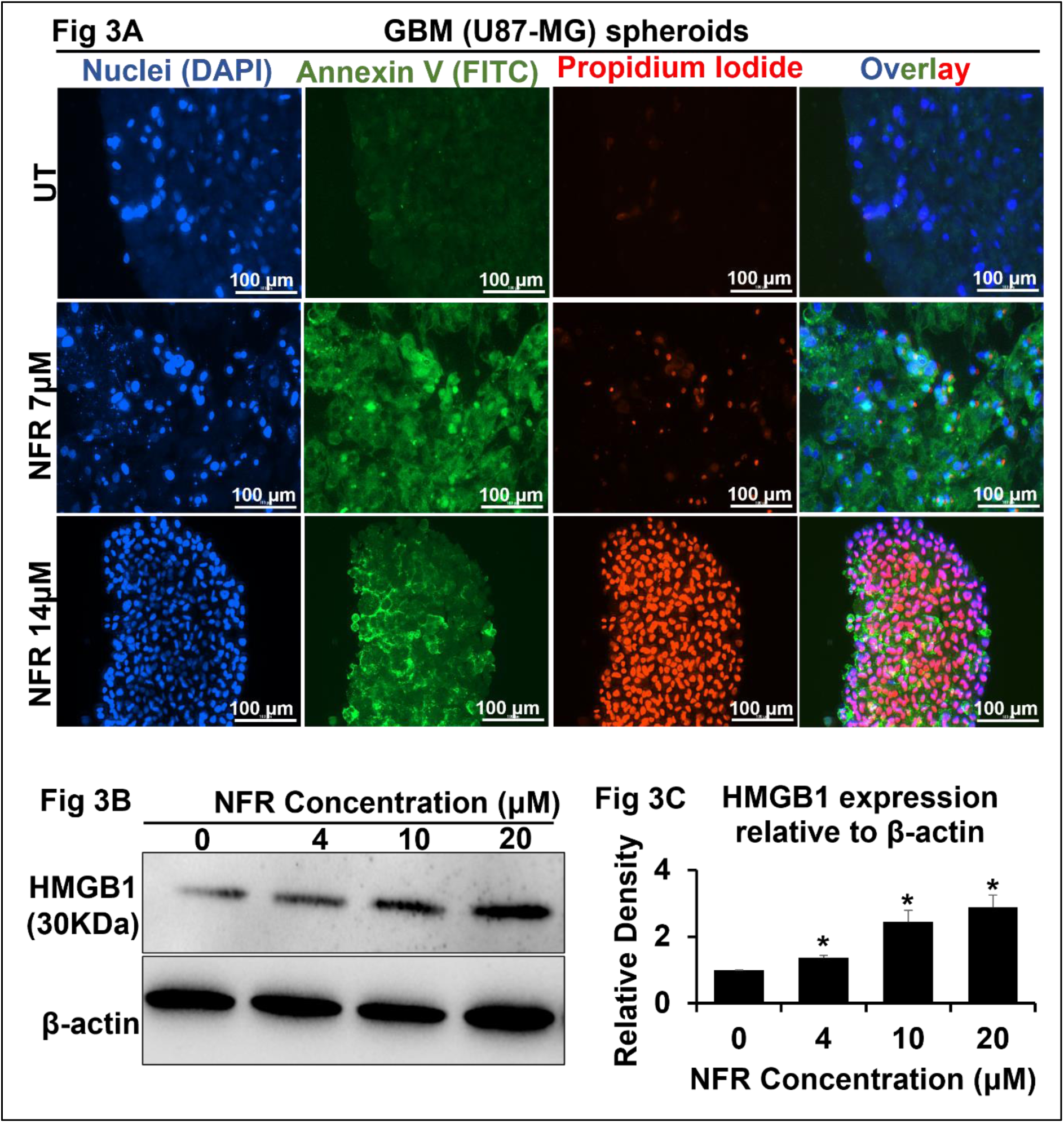
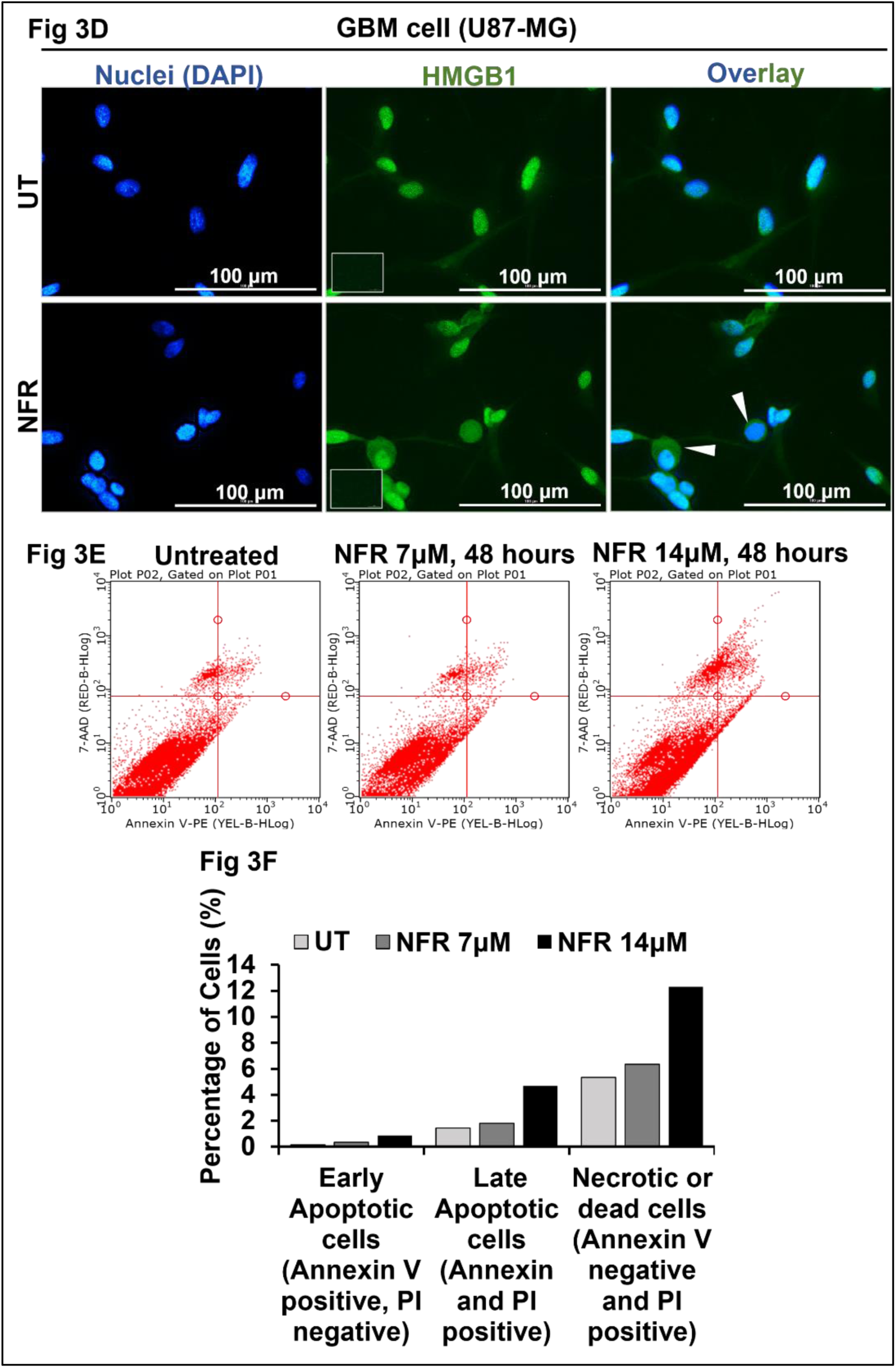
Nelfinavir induces Apoptosis and Necrosis in Glioblastoma cells: A. UT and NFR (7μM, 14μM, 48 hours) treated GBM (U87-MG) spheroids were stained with annexin V (green), propidium iodide (PI) (red), and DAPI (nuclei, blue). At least 3 frames were imaged per spheroid. Images are representative of all frames. Scale bar,100 µm. B. HMGB1 expression is quantified in NFR (4μM, 10μM, 20μM, 24 hours) treated GBM (U87-MG) cells using western blot. β-actin is used as a loading control. 15µg protein sample is loaded in each well. Images are representative of 3 experiments. C. The western blot data were quantified using densitometry. The graph is representative of 2 experiments. D. UT and NFR (10μM, 24 hours) treated GBM (U87-MG) cells were stained with anti-HMGB1 antibody (green) and DAPI (nuclei, blue). At least 10 frames were imaged per well of cells cultured in a well chamber slide. Images are representative of all frames. The inset represents a primary antibody control. In overlay, the inset represents the zoomed-in area of the image. Scale bar,100 µm. E. GBM (U87-MG) spheroids were treated with NFR (7μM, 14μM, 48 hours), and spheroids were subsequently stained with FITC-conjugated annexin V and propidium iodide (PI) and analyzed by flow cytometry. F. Flow cytometry data was quantified. The graph represents the percentage of cells undergoing apoptosis and necrosis in NFR-treated GBM (U87-MG) spheroids. Data are represented as mean ± SEM. Error bars indicate SEM. * p < 0.05, **p < 0.01, ***p < 0.005 (Student’s t-test).

### 4. Nelfinavir promotes AIM2 inflammasome activation in Glioblastoma cells

Previous studies in monocytes (THP-1) have shown that NFR activates the AIM2 inflammasome by promoting the release of nuclear DNA into the cytoplasm, due to impaired nuclear membrane maturation [17]. In our study with GBM cells, we observe increased expression of HMGB1 following NFR treatment (Figure 3B, 3C). Notably, increased HMGB1 expression is a known marker of inflammation, and it has also been reported to support AIM2 inflammasome activation in hepatocytes by binding with the HIN domain of AIM2 [48,49]. Since HMGB1 remains bound to DNA, it functions as a chaperone that enhances dsDNA recognition by AIM2. Based on these observations, we investigated whether NFR similarly triggers AIM2 inflammasome activation in GBM cells. To determine whether NFR compromises the nuclear membrane in GBM cells, as previously observed in THP-1 cells, we treated GBM cells with NFR and performed immunocytochemistry using Anti-Lamin B1 antibody and nuclear DNA staining with DAPI (Figure 4A). Lamin B1 is a key structural protein of the nuclear lamina that plays vital roles in maintaining nuclear structure and integrity [50]. Lamin B1 is synthesized as a precursor protein that undergoes post-translational modifications to become fully functional and properly localized. Impaired Lamin B1 maturation or decreased Lamin B1 expression can lead to nuclear membrane instability and subsequent release of nuclear DNA. As shown in Figure 4A, NFR-treated GBM cells exhibited the release of nuclear DNA into the cytoplasm, along with disrupted Lamin B1 expression. To validate these findings, we conducted a western blot for Lamin B1, which revealed a decreased expression of Lamin B1 following NFR treatment (Figure 4B, C). This data demonstrates that NFR disrupts the nuclear membrane by disrupting the Lamin B1 expression in GBM cells, leading to the release of nuclear DNA into the cytoplasm. These findings are consistent with previously published data in THP-1 cells, where NFR was shown to induce nuclear DNA release, a known trigger for AIM2 inflammasome activation [17].

**Legend Figure 4.**
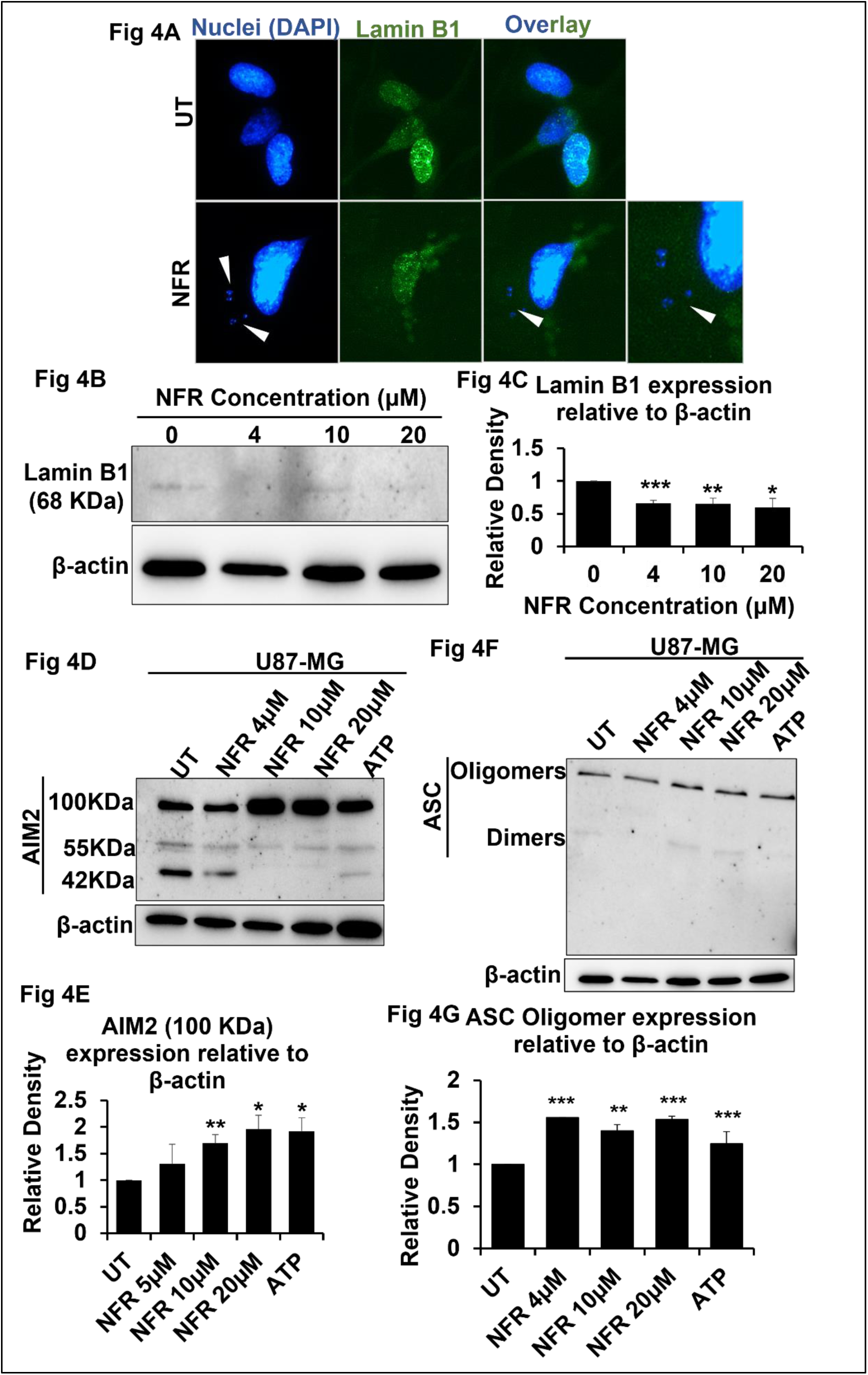
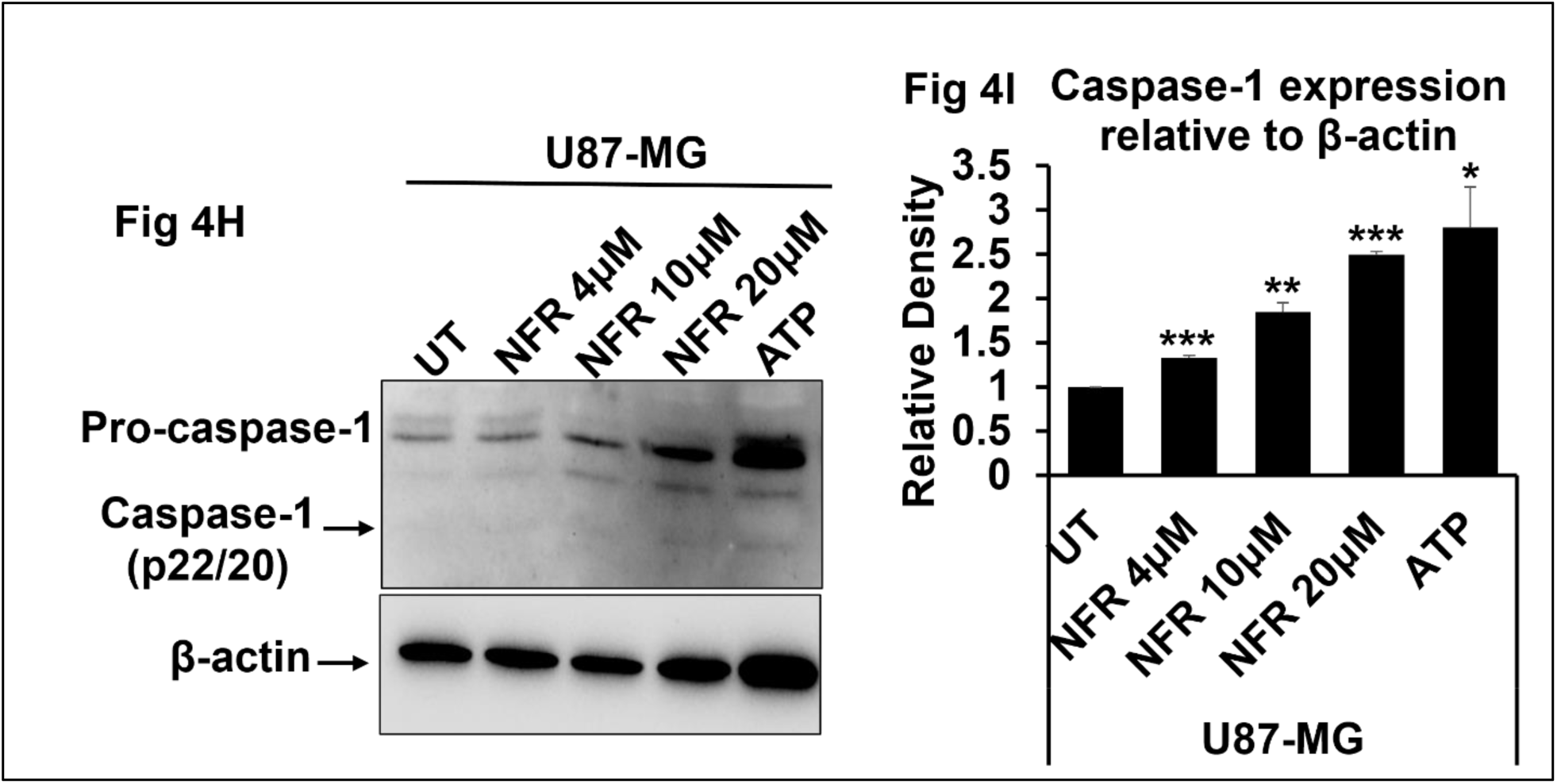
Nelfinavir promotes AIM2 inflammasome activation in Glioblastoma cells: A. To check the Lamin B1 expression in UT and NFR (10μM, 24 hours) treated GBM (U87-MG) cells, they were stained with anti-Lamin B1 antibody (green) and DAPI (nuclei, blue). At least 10 frames were imaged per well of the two-well chamber slides. Images are representative of all frames. B. Lamin B1 expression is quantified in NFR (4μM, 10μM, 20μM, 24 hours) treated GBM (U87-MG) cells using western blot. β-actin is used as a loading control. 15µg protein sample is loaded in each well. Images are representative of 3 experiments. C. The western blot data was quantified using densitometry. The graph is representative of 3 experiments. D, F, H. AIM2, ASC, and Caspase-1 expression is quantified in LPS (0.5 μg/ml, 6 hours) primed NFR (4μM, 10μM, 20μM, 24 hours) treated GBM (U87-MG) cells using western blot. β-actin is used as a loading control. 15µg protein sample is loaded in each well. Images are representative of 2 experiments. E, G, I. The western blot data was quantified using densitometry. The graph is representative of two experiments. Data are represented as mean ± SEM. Error bars indicate SEM. * p < 0.05, **p < 0.01, ***p < 0.005 (Student’s t-test).

Next, we assessed whether NFR-induced nuclear DNA release could trigger AIM2 activation in GBM cells. We used LPS-primed and ATP-treated GBM cells, along with LPS-primed THP-1 cells, as positive controls for inflammasome activation [51]. The western for AIM2 in NFR-treated GBM cells shows three distinct bands, with molecular weights of approximately 40 kDa, 55 kDa, and ∼100 kDa. The 40 kDa and 55 kDa bands represent monomeric forms of AIM2, corresponding to its two known isoforms [52] (Figure 4D, E). The ∼100 kDa band likely represents the oligomeric form of AIM2, which forms upon inflammasome assembly with ASC (Apoptosis-associated Speck-like protein containing a CARD) and pro-caspase-1 [53]. Following NFR treatment, we observed an increase in the ∼100 kDa oligomeric AIM2 band (Figure 4E) and a concomitant decrease in the 40 kDa monomeric AIM2 band. This suggests that monomeric AIM2 is recruited into inflammasome complexes, leading to the formation of the higher molecular weight oligomeric structure[54]. To further validate inflammasome activation, we examined ASC oligomerization and pro-caspase-1 cleavage in NFR-treated cells using western blot analysis (Figures 4F, G, H, I). As shown in Figures 4F and 4G, NFR treatment led to increased levels of oligomerized ASC, indicating ASC polymerization. Additionally, Figures 4H-I show a significant increase in the cleaved (p20) form of caspase-1 following NFR treatment, confirming caspase-1 activation. Together, these results confirm that NFR treatment induces AIM2 inflammasome activation in GBM cells. These findings are consistent with previously published data in THP-1 cells, where NFR was shown to induce AIM2 inflammasome activation, leading to caspase-1 activation and ASC oligomerization [17].

### 5. Differential expression of AIM2 in Glioma patients and effect of Nelfinavir on patient-derived glioma cells and organoids

We next investigated AIM2 expression in diverse GBM cell lines, glioma patient samples, and assessed the impact of NFR on the viability of patient-derived primary glioma cells and organoids. We examined the expression of AIM2 in various GBM cell lines, including LN-18, LN-229, and U87-MG, as well as in the astrocyte cell line SVG (Figure S4A-C). The expression pattern of AIM2 exhibits variability among different cells, as depicted in Figure S4A. Specifically, AIM2 expression in U87-MG cells exhibits a distinct punctate pattern, whereas other cell types (LN-229, LN-18, SVG) display a more diffuse AIM2 distribution. The variation in expression pattern may be attributed to distinct cell-type-specific regulation or function of AIM2 signaling pathways. The LN-229 and LN-18 glioma cell lines harbor mutant TP53 and likely possess homozygous deletions in the tumor suppressor genes *CDKN2A* (cyclin-dependent kinase inhibitor 2A). In contrast, the U87-MG cell line is characterized by a hypodiploid karyotype, with a modal chromosome number of 44 observed in approximately 48% of the cells. Further, to quantify the expression of AIM2 in different types of cells, western blot analyses were performed utilizing cell lysates from GBM and astrocyte cell lines (Figure S4B). Densitometry analyses indicate variation in AIM2 total baseline protein expression across GBM cell types (Figure S4C). Further, to evaluate AIM2 expression in glioma patients (grades 2 and 3), we collected matched normal and tumor brain tissues from glioma patients (Table 1). Western blot analysis of tissue lysates revealed a consistent decrease in AIM2 expression in all glioma samples compared to their matched normal controls (Figure 5A–C). AIM2 was detected as a doublet at ∼55 kDa and ∼40 kDa (Figure 5A) in all patient samples. This is similar to previous reports with GBM tumor tissue[52]. We also performed western blot analysis for caspase-1 expression in glioma tissue samples obtained from three different patients (Figure S4D-E). Notably, AIM2 and Caspase-1 expression varied across patients, potentially reflecting differences in glioma grade and the heterogeneity of the tumor microenvironment (TME). Glioma is a highly heterogeneous tumor characterized by a complex TME composed of malignant cells, immune cells, vascular components, stem-like cells, and various glial cell types [55]. Among the glial components within the glioma TME, astrocytes contribute to tumor progression by secreting neurotrophic factors such as growth differentiation factor 15 (GDF-15) and glial cell line-derived neurotrophic factor (GDNF), which promote GBM cell proliferation and migration, respectively [56,57]. Microglia, the resident immune cells of the central nervous system (CNS), also play a crucial role in gliomagenesis by enhancing glioma cell invasion and supporting tumor growth [58].

**Legend Figure 5.**
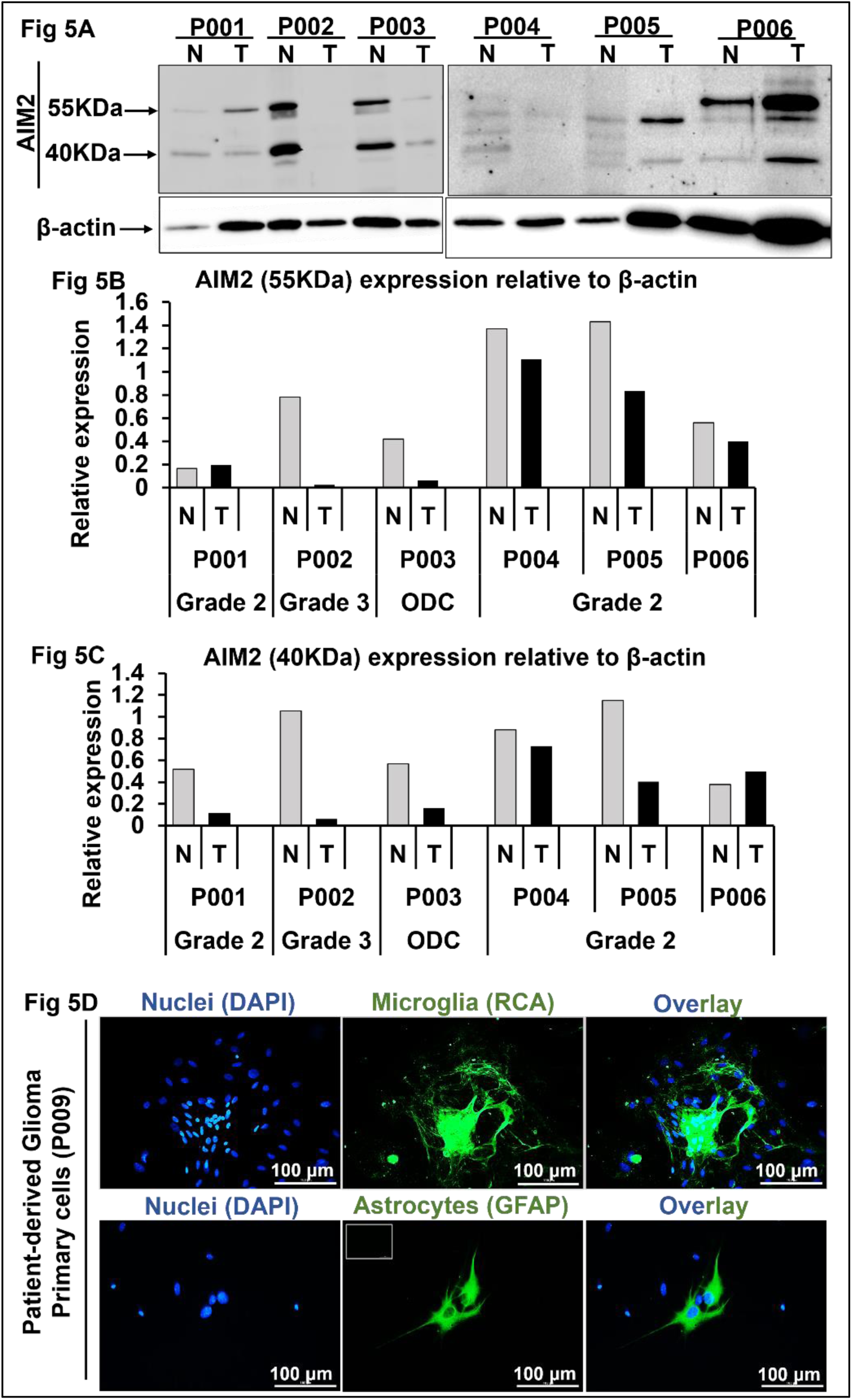
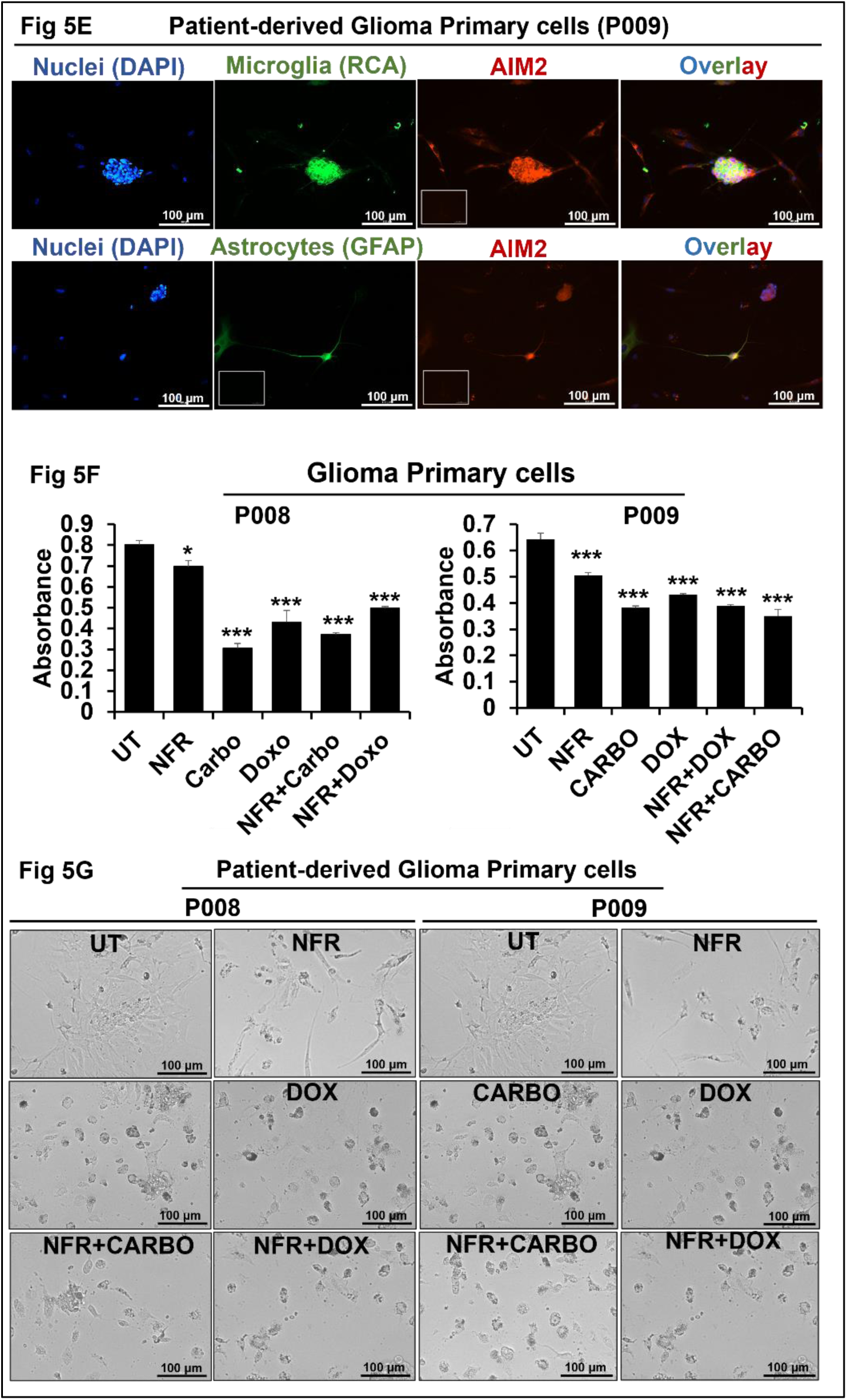
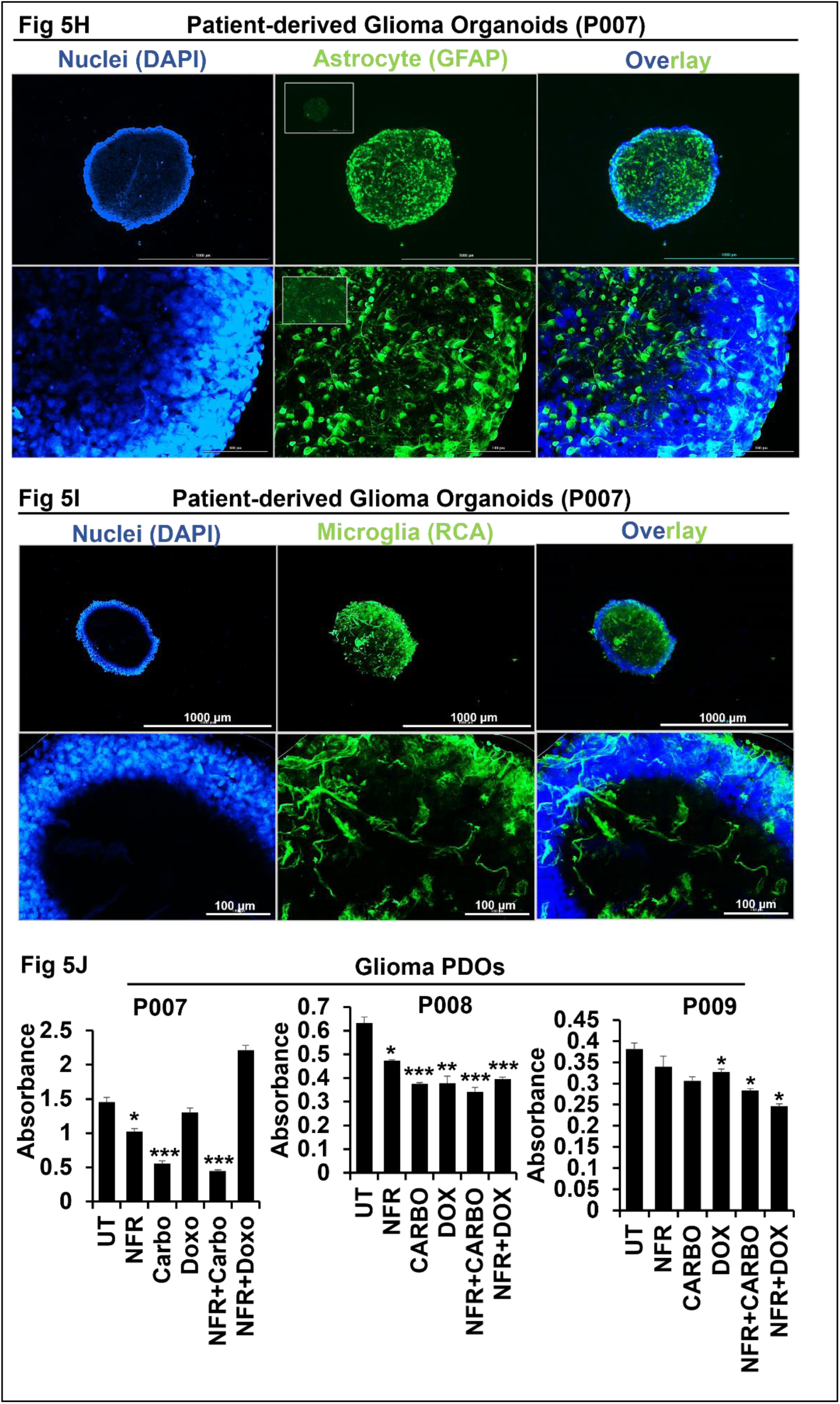
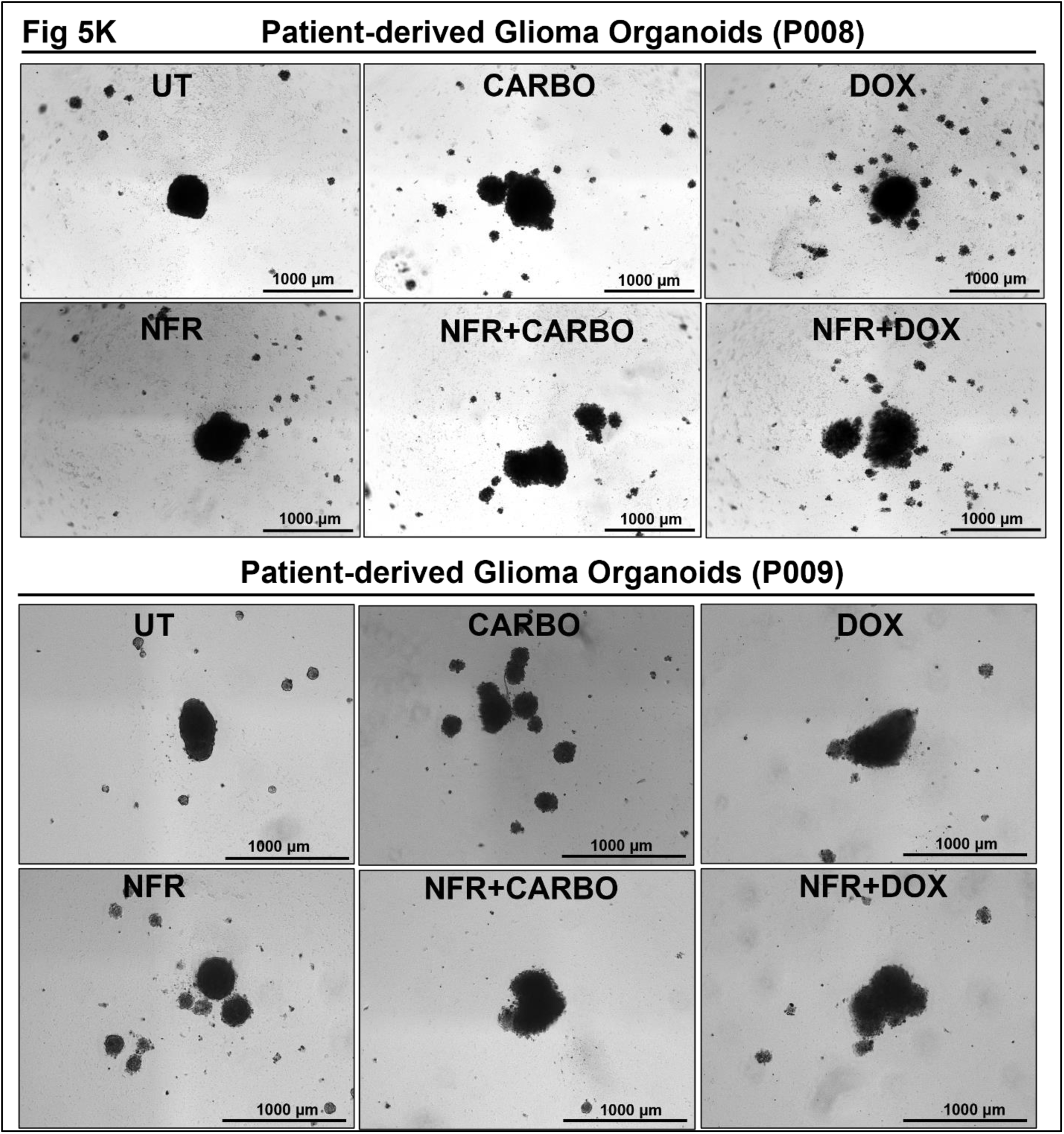
Differential expression of AIM2 in Glioma patients and effect of Nelfinavir on Glioma patient-derived cells and organoids: A. AIM2 expression was quantified across diverse glioma-grade patient tissues using western blot. β-actin is used as a loading control (N = normal, T = tumor, ODC = other disease control) (n=6). B, C. The western blot data was quantified using densitometry. D. Patient-derived primary glioma cells (P009) were stained with anti-GFAP antibody and RCA to detect astrocytes and microglia, respectively. DAPI was used to stain nuclei (blue). At least 7 frames were imaged per section. Images are representative of all frames. Inset represents primary antibody control. Scale bar, 100μm. E. Patient-derived primary glioma cells (P009) were stained with anti-AIM2 antibody, anti-GFAP, and RCA. The co-localization of AIM2 with astrocytes and microglia can be visualized by the red staining overlapping with green. DAPI was used to stain nuclei (blue). At least 7 frames were imaged per section. Images are representative of all frames. Inset represents primary antibody control. Scale bar, 100μm. F. Cell viability assay was performed in patient-derived primary glioma cells (P008, P009) treated with NFR (4μM), Carboplatin (CARBO, 2.7mM), Doxorubicin (DOX, 4.9μM), or in indicated combination for 48 hours. The graph is representative of 3 replicates. G. Brightfield images of patient-derived primary glioma cells (P008, P009) treated with NFR, CARBO, DOX, or in indicated combination for 48 hours. At least 7 frames were imaged per well. Images are representative of all frames. Scale bar, 100μm. H. Patient-derived glioma organoids (P007) were stained with anti-GFAP antibody to detect astrocytes. DAPI was used to stain nuclei (blue). Inset represents primary antibody control. Scale bar, 1000μm, 100μm. I. Patient-derived glioma organoids (P007) were stained with RCA to detect microglia. DAPI was used to stain nuclei (blue). Scale bar, 1000μm, 100μm. J. Cell viability assay was performed in patient-derived glioma organoids (P007, P008, P009) treated with NFR (7μM), CARBO (2.7mM), DOX (4.9μM), or in indicated combination for 48 hours. The graph is representative of 3 replicates. K. Brightfield images of patient-derived glioma organoids (P008, P009) treated with NFR, CARBO, DOX, or in indicated combination for 48 hours. Scale bar, 1000μm. Data are represented as mean ± SEM. Error bars indicate SEM. * p < 0.05, **p < 0.01, ***p < 0.005 (Student’s t-test). PDO=patient-derived organoid.

**Table 1.** Details of patients: This table summarizes the clinical details of glioma patients used in the study, including sex, age at diagnosis, tumor type, WHO grade, and key findings from histopathological evaluation. M: male; F: female, CNS: central Nervous system, WHO: World Health Organisation.

| Patient number | Age/Sex | Pathology report |
| --- | --- | --- |
| P001 | 22/M | Central Neurocytoma (CNS WHO Grade 2) |
| P002 | 45/F | Astrocytoma (CNS WHO Grade 3) |
| P003 | 49/F | Neuroendocrine Carcinoma |
| P004 | 48/M | Diffuse astrocytoma with gemistocytic and oligodendroglial areas (CNS WHO Grade 2) |
| P005 | 32/M | Oligodendroglioma (CNS WHO Grade 2) |
| P006 | 30/M | Diffuse Astrocytoma with oligodendroglial (CNS WHO Grade 2) |
| P007 | 39/M | Oligodendroglioma (CNS WHO grade 2) |
| P008 | 16/F | Anaplastic pleomorphic xanthoastrocytoma (CNS WHO grade 3) |
| P009 | 28/F | Pilocytic astrocytoma (CNS WHO grade 1) |

To evaluate the effect of NFR on a more clinically relevant model, we isolated primary glioma cells from freshly resected patient tumor tissues (Table 1). To assess the cellular heterogeneity of these primary glioma cultures, we evaluated the presence of astrocytes and microglia, as both cell types represent major components of the glioma TME [59]. Fluorescence microscopy was performed using glial fibrillary acidic protein (GFAP) as a marker for astrocytes and Ricinus communis agglutinin (RCA) for microglia. As shown in Figure 5D, both astrocytes and microglia were readily detected in the patient-derived primary glioma cells, confirming cellular heterogeneity. Furthermore, immunostaining in Figure 5E demonstrated the presence of AIM2 in both microglia and astrocytes within these cultures, suggesting that AIM2 is expressed across multiple cell types in the glioma TME. Subsequently, we treated these patient-derived primary glioma cells with NFR (4μM), Carboplatin (2.7mM), Doxorubicin (4.9μM), and combinations of NFR + Carboplatin and NFR + Doxorubicin. MTT assay results revealed a significant reduction in cell viability upon NFR treatment, either alone or in combination (Figure 5F). In Figure 5G, the images of primary glioma cells treated with drugs further confirm our MTT results. These findings indicate that NFR is effective not only in established glioma cell lines but also in glioma cells derived from patients.

Several models have been developed to study glioma growth dynamics and to investigate treatment-related, cell-intrinsic pathways and mechanisms [60–63]. However, conventional models often fail to replicate the complexity of glioma stem cell (GSC) heterogeneity, cell–cell interactions, and treatment responses, representing a major limitation of these models. Patient-derived xenograft (PDX) models, created by implanting freshly resected tumor tissue into the brains of immunocompromised mice, are widely regarded as more accurate in recapitulating tumor heterogeneity compared to traditional models [64,65]. Nevertheless, PDX models are hindered by challenges such as long latency periods, genetic drift, and epigenetic alterations that can compromise their clinical relevance [66]. More recently, the development of patient-derived organoids (PDOs) has emerged as a promising preclinical platform. Glioma PDOs have been shown to preserve key histological, cellular, genomic, and transcriptomic characteristics of the original tumors, and can be generated within a clinically relevant timeframe to facilitate personalized therapeutic testing [67,68]. To evaluate the therapeutic efficacy of NFR in a clinically relevant setting, we generated glioma organoids from freshly resected, patient-derived tumor tissues (Table 1). To characterize the cellular composition of these organoids, we performed immunohistochemistry analyses. Astrocytes were identified with Anti-GFAP, and microglia were identified with RCA staining. Figures 5H-I demonstrate the presence of astrocytes and microglia within organoids. These cell types are critical components of the glioma TME, known to influence tumor progression and therapeutic response [59]. Astrocytes contribute to tumor progression by secreting neurotrophic factors that enhance GBM cell proliferation and migration [56,57]. Moreover, the proportion of microglia and macrophages in GBM correlates with the histological grade of gliomas, and these cells exhibit immunosuppressive and pro-invasive properties that facilitate tumor progression [59]. The preservation of these populations in the organoid model underscores its fidelity to the parental tumor and highlights its potential as a robust platform for patient-specific drug testing and mechanistic studies. Glioma patient-derived organoids were subsequently treated with NFR (7μM), Carboplatin (2.7mM), Doxorubicin (4.9μM), both individually and in combination, followed by a cell viability assay to evaluate therapeutic efficacy. As shown in Figure 5J drug treatment led to a reduction in cell viability, with NFR demonstrating a variable degree of cytotoxicity across different glioma patient-derived organoids. Figure 5K presents representative images of glioma organoids following drug treatment. The drug-treated organoids exhibit a disrupted and loosened spheroid architecture, suggestive of drug-induced cell death. Notably, NFR was more effective in organoids derived from higher-grade gliomas (grades 2 and 3) compared to those from lower-grade tumors (grade 1). This differential response likely reflects inter-and intratumoral heterogeneity, underscoring the importance of developing precision or personalized therapeutic strategies tailored to the specific characteristics of each patient’s tumor.

## Discussion and Conclusion

The anti-tumor properties of HIV protease inhibitor NFR have been observed in diverse cancer types, including breast, melanoma, prostate, and non-small lung cancers [11,13,14,20]. In this study, we demonstrate the anti-tumor efficacy of the NFR in human GBM cells. We observed that NFR effectively inhibits GBM cell growth with an IC₅₀ of ∼5μM (Figure 1A-B). *In vitro* assays revealed that NFR inhibits GBM cell proliferation and induces both apoptotic and necrotic cell death (Figure 1C-D, 3A-F). These findings suggest that the anti-tumor activity of NFR in GBM cells is linked to its anti-proliferative and pro-apoptotic/necrotic properties. Previous studies have reported the anti-proliferative properties of NFR in various cancers, including breast, prostate, and glioma [11,13,33]. Moreover, NFR has been shown to induce apoptosis and/or necrosis in melanoma, breast cancer, and NSCLC cell lines [11,14,16]. The molecular mechanisms underlying NFR’s anti-cancer effects are diverse, involving alterations in androgen receptor (AR), STAT3, and Akt signaling pathways, as well as cell cycle arrest. In GBM specifically, NFR has been shown to trigger endoplasmic reticulum stress and inhibit angiogenesis [15,33]. Another interesting finding from our study states that NFR activates the AIM2 inflammasome in GBM cells by inducing the release of nuclear DNA into the cytoplasm (Figure 4A-H). AIM2 inflammasome activation can act as an additional mechanism that may contribute to the anti-tumor effects of NFR in GBM cells. AIM2 is a member of the interferon-inducible p200 protein family and plays a key role in the innate immune response against both exogenous and endogenous pathogens, including *Francisella tularensis*, *Listeria monocytogenes*, and various DNA viruses [69,70]. More recent studies have identified AIM2 as a novel pattern recognition receptor (PRR) that detects cytosolic double-stranded DNA (dsDNA) [71]. Upon recognizing dsDNA, AIM2 recruits the adaptor protein ASC to assemble a caspase–1–activating inflammasome complex [72]. AIM2 recognizes dsDNA through its C-terminal HIN-200 domain, which contains oligonucleotide/oligosaccharide-binding folds, while its N-terminal pyrin domain is responsible for recruiting ASC [73,74]. Activation of the AIM2 inflammasome leads to the cleavage of pro–IL–1β and pro–IL–18, resulting in the release of their biologically active forms, IL-1β and IL-18 [53]. While the role of AIM2 in innate immunity and inflammation is well established, its function in cancer remains complex and context-dependent. AIM2 has been reported to exhibit dual roles in tumor development [75]. Initially, AIM2 was identified as a tumor suppressor gene [76]. Loss of AIM2 has been associated with enhanced metastasis in hepatocellular carcinoma through inflammasome-dependent pathways, and its overexpression has been shown to inhibit migration and invasion of renal carcinoma cells [77,78]. However, recent studies suggest that AIM2 may also act as an oncogene in certain cancer types. For example, in NSCLC, oral squamous cell carcinoma, and Epstein-Barr virus (EBV)–associated nasopharyngeal carcinoma, AIM2 has been implicated in promoting tumor progression through inflammasome activation [79–81]. Additionally, AIM2 has been shown to exert tumor-suppressive effects independent of inflammasome activation, particularly in colorectal cancer [82]. In the context of GBM, earlier research indicates that AIM2 can suppress GBM cell proliferation [52]. Therefore, it is plausible that one of the mechanisms by which NFR exerts its anti-proliferative effects in GBM cells is through the activation of AIM2. In line with our functional assays, western blot analysis of patient-derived tissue samples revealed reduced AIM2 expression in glioma tissues compared to adjacent normal brain tissues (Figure 5A-C). This observation supports our hypothesis that AIM2 may function as a tumor suppressor in GBM. Previous studies have demonstrated that AIM2 expression is frequently downregulated in several cancers, including renal cancer and hepatocellular carcinoma, where its loss promotes tumor progression and immune evasion through suppression of inflammasome activity [77,78]. Our findings align with these reports and highlight a potential mechanistic link between AIM2 suppression and glioma pathobiology. Together, this suggests that AIM2 downregulation may contribute to glioma progression and that restoration of AIM2 activity could offer a novel therapeutic angle.

Conventional cell culture on plastic dishes is often considered suboptimal for evaluating cancer therapeutics, as it fails to mimic the interactions between cancer cells accurately. To better assess the efficacy of NFR as a tumor suppressor, we utilized a 3D spheroid model in which GBM cells were grown as spheroids. 3D models can more closely approximate *in vivo* tumor behaviour and therefore offer improved predictive value for evaluating novel cancer therapies. In this assay, NFR demonstrated strong anti-tumor activity, effectively inhibiting the growth of GBM cells (Figure 1E-M). The effective NFR concentration in the GBM 3D model was 7μM, higher than that required in 2D culture (∼5μM), likely reflecting the enhanced capacity of cancer cells to thrive in 3D environments that more closely resemble *in vivo* conditions. We observed that the IC₅₀ value of NFR was substantially higher in astrocyte spheroids compared to GBM spheroids. These findings contrast with our 2D culture data, where the IC₅₀ values of NFR were nearly identical across both cell types (Figure 1A-B). This differential sensitivity may be attributed to the distinct microenvironmental context provided by 3D culture systems. Prior studies have shown that astrocytes cultured in 3D environments recapitulate key morphological and biochemical features of *in vivo* astrocytes, which are typically lost in conventional 2D culture [83]. The 3D architecture supports physiologically relevant filopodial dynamics and significantly attenuates the cellular stress commonly associated with 2D conditions. Additionally, astrocytes maintained in 3D microenvironments more accurately reflect *in vivo* phenotypes in terms of astrocyte–endothelial interactions and activation status, as indicated by GFAP expression [84]. Thus, the astrocytes cultured in a 3D microenvironment more closely reflect the *in vivo* response to a drug. Notably, the concentrations used for our study are comparable to the peak plasma levels observed in patients and healthy volunteers receiving the recommended dosage of NFR. In a phase I/II clinical trial, HIV-infected patients were treated with 500 or 750 mg of NFR three times daily, resulting in peak plasma concentrations of 3 to 4 μg/mL (equivalent to 4.5–6.0μmol/L) [31]. We further evaluated the efficacy of NFR in comparison with standard chemotherapeutic agents, Doxorubicin and Carboplatin. NFR exhibited synergistic cytotoxic effects when combined with Doxorubicin or Carboplatin in both 2D and 3D GBM cell culture models (Figure 2). Several Phase I clinical trials across a range of cancer types, including locally advanced cervical cancer, stage IIIA/IIIB NSCLC, locally advanced pancreatic cancer, multiple myeloma, and GBM, have demonstrated that NFR, when combined with standard chemotherapies (cisplatin, bortezomib, TMZ) along with radiotherapy, is both feasible and well-tolerated [21,85–88]. Our study provides additional insights into the potential of combinatorial therapeutic strategies for GBM.

While spheroid models have been instrumental in advancing our understanding of GBM biology, organoid models offer several distinct advantages that make them increasingly preferable for translational research [89]. Spheroids are primarily formed through the aggregation of established cell lines, whereas organoids are typically derived from primary cells, including patient tumor tissues. GBM organoids more accurately recapitulate the tumor’s architectural complexity, cellular heterogeneity, and microenvironmental interactions, including the presence of hypoxic gradients and stem cell niches, which are often absent or poorly defined in spheroids [60,61,63,67]. Furthermore, organoids derived from patient tumor tissues retain key genetic, epigenetic, and phenotypic features of the original tumor, making them a more physiologically relevant platform for studying disease progression and therapeutic responses [90,91]. Organoid-based models represent a powerful tool for preclinical drug screening and personalized medicine in GBM. To evaluate therapeutic efficacy in a more physiologically relevant system, we generated organoids from glioma patient tissue samples (WHO Grade 1–3) and treated them with NFR, Carboplatin, and Doxorubicin (Figure 5H-K). Notably, we observed heterogeneous drug responses across the patient-derived glioma organoids, which can be attributed, in part, to the inherent molecular and phenotypic diversity associated with glioma grades (WHO Grade 1–3) [1]. While all these tumors (Grade 1–3) were classified as lower-grade gliomas, they differ considerably in terms of growth kinetics, differentiation status, and underlying genetic alterations. Grade 1 gliomas, such as pilocytic astrocytomas, are generally slow-growing and frequently harbor MAPK pathway alterations, including BRAF mutations [92]. In contrast, Grade 2 and 3 gliomas are more aggressive and commonly exhibit IDH1/2 mutations, ATRX loss, and in some cases, 1p/19q co-deletion, genetic changes that influence tumor metabolism, DNA repair capacity, and treatment sensitivity [93]. These molecular distinctions may affect not only how cells respond to cytotoxic stress but also how effectively drugs are taken up and retained [94]. Additionally, variability in tumor cell composition, particularly in the proportion of glioma stem-like cells, and structural differences between organoids, such as density and size, may influence both drug diffusion and therapeutic outcome [61,67].

Our findings demonstrate that NFR possesses significant anti-tumor activity in GBM, mediated through both anti-proliferative and cell death–inducing mechanisms. Using a comprehensive range of GBM models, including established cell lines, cell-derived spheroids, patient-derived primary glioma cells, and organoids, we showed that NFR effectively reduces cell viability and induces both apoptotic and necrotic cell death. Moreover, NFR exhibited synergistic effects when combined with conventional chemotherapeutic agents, Carboplatin and Doxorubicin, across these models, suggesting its potential utility in combination therapy. Mechanistically, we observed activation of the AIM2 inflammasome following NFR treatment, implicating this pathway as a possible contributor to its anti-tumor effects. Together, these results highlight NFR as a promising therapeutic candidate for GBM and point toward AIM2 inflammasome activation as a novel mechanism warranting extended Phase I clinical trial studies investigating the combination of NFR, temozolomide, and radiotherapy in patients with GBM.

## Supporting information

supplementray figures with legend

## Statements and Declarations

## Acknowledgment

S.J.’s laboratory was founded with institutional funds from the Indian Institute of Technology Jodhpur (IITJ) and the generation of patient-derived spheroids in this study was funded by the Ministry of Electronics and Information Technology, Government of India (No.4 (16)/2019-ITEA). Dr. Pankaj Seth from the National Brain Research Centre (NBRC) has provided the human astrocyte cell line SVG. Sections of human brain tissue were procured from the All-India Institute of Medical Sciences (AIIMS) Jodhpur and Tata Memorial Hospital, Tata Memorial Center, Mumbai, Maharashtra, India. We express our gratitude to Mr. Bharat Pareek, Technical Superintendent at the Indian Institute of Technology Jodhpur, for his technical assistance in our laboratory.

## Data Availability Statement

The data generated in this study are available within the article and its supplementary data files.

## Ethical Approval

Fresh and Paraffin-embedded, paraformaldehyde-fixed glioma and normal brain tissue were acquired with authorization from the Internal Review Board and the Ethics Committees of the All-India Institute of Medical Sciences (AIIMS), Jodhpur, and Tata Memorial Cancer Hospitals. All studies were conducted in compliance with the ethical rules and regulations of the All-India Institute of Medical Sciences (AIIMS), Jodhpur, and the Indian Institute of Technology Jodhpur.

## Consent to participate

Informed consent was obtained from all individual participants included in the study.

## Consent for publication

Informed consent was obtained from all individual participants included in the study.

## Author Contribution

D.M. designed and performed the experiments (data analysis, immunofluorescence, western blotting, primary cell isolation) and prepared and edited the initial manuscript draft. S.C. and P.S. did CFA quantification. L.S. helped with flow cytometry experiments. S.J. conceptualized the study, designed experiments, and edited and reviewed the manuscript. All authors reviewed the manuscript.

## References

[1] Louis DN, Perry A, Wesseling P, Brat DJ, Cree IA, Figarella-Branger D, et al. The 2021 WHO classification of tumors of the central nervous system: A summary. Neuro Oncol 2021;23:1231–51. 10.1093/neuonc/noab106.

[2] Aldape K, Zadeh G, Mansouri S, Reifenberger G, von Deimling A. Glioblastoma: pathology, molecular mechanisms and markers. Acta Neuropathol 2015;129:829–48. 10.1007/s00401-015-1432-1.

[3] Hotchkiss KM, Karschnia P, Schreck KC, Geurts M, Cloughesy TF, Huse J, et al. A brave new framework for glioma drug development. Lancet Oncol 2024;25:e512–9. 10.1016/S1470-2045(24)00190-6.

[4] Horbinski C, Nabors LB, Portnow J, Baehring J, Bhatia A, Bloch O, et al. NCCN GUIDELINES® INSIGHTS: Central Nervous System Cancers, Version 2.2022. JNCCN Journal of the National Comprehensive Cancer Network 2023;21:12–20. 10.6004/jnccn.2023.0002.

[5] Stupp R, Mason WP, Van Den Bent MJ, Weller M, Fisher B, Taphoorn MJB, et al. Radiotherapy plus Concomitant and Adjuvant Temozolomide for Glioblastoma. n.d.

[6] Stupp R, Taillibert S, Kanner A, Read W, Steinberg DM, Lhermitte B, et al. Effect of tumor-treating fields plus maintenance temozolomide vs maintenance temozolomide alone on survival in patients with glioblastoma a randomized clinical trial. JAMA - Journal of the American Medical Association 2017;318:2306–16. 10.1001/jama.2017.18718.

[7] Subeha MR, Telleria CM. The anti-cancer properties of the hiv protease inhibitor nelfinavir. Cancers (Basel) 2020;12:1–38. 10.3390/cancers12113437.

[8] Rascio F, Spadaccino F, Rocchetti MT, Castellano G, Stallone G, Netti GS, et al. The pathogenic role of PI3K/AKT pathway in cancer onset and drug resistance: an updated review. Cancers (Basel) 2021;13. 10.3390/cancers13163949.

[9] Subeha MR, Telleria CM. The anti-cancer properties of the hiv protease inhibitor nelfinavir. Cancers (Basel) 2020;12:1–38. 10.3390/cancers12113437.

[10] Pyrko P, Kardosh A, Wang W, Xiong W, Schönthal AH, Chen TC. HIV-1 protease inhibitors nelfinavir and atazanavir induce malignant glioma death by triggering endoplasmic reticulum stress. Cancer Res 2007;67:10920–8. 10.1158/0008-5472.CAN-07-0796.

[11] Soprano M, Sorriento D, Rusciano MR, Maione AS, Limite G, Forestieri P, et al. Oxidative stress mediates the antiproliferative effects of nelfinavir in breast cancer cells. PLoS One 2016;11. 10.1371/journal.pone.0155970.

[12] Gupta AK, Cerniglia GJ, Mick R, McKenna WG, Muschel RJ. HIV protease inhibitors block Akt signaling and radiosensitize tumor cells both in vitro and in vivo. Cancer Res 2005;65:8256–65. 10.1158/0008-5472.CAN-05-1220.

[13] Yang Y, Ikezoe T, Takeuchi T, Adachi Y, Ohtsuki Y, Takeuchi S, et al. HIV-1 protease inhibitor induces growth arrest and apoptosis of human prostate cancer LNCaP cells in vitro and in vivo in conjunction with blockade of androgen receptor STAT3 and AKT signaling. Cancer Sci 2005;96:425–33. 10.1111/j.1349-7006.2005.00063.x.

[14] Jiang W, Mikochik PJ, Ra JH, Lei H, Flaherty KT, Winkler JD, et al. HIV protease inhibitor nelfinavir inhibits growth of human melanoma cells by induction of cell cycle arrest. Cancer Res 2007;67:1221–7. 10.1158/0008-5472.CAN-06-3377.

[15] Pore N, Gupta AK, Cerniglia GJ, Maity A. HIV protease inhibitors decrease VEGF/HIF-1α expression and angiogenesis in glioblastoma cells. Neoplasia 2006;8:889–95. 10.1593/neo.06535.

[16] Yang Y, Ikezoe T, Nishioka C, Bandobashi K, Takeuchi T, Adachi Y, et al. NFV, an HIV-1 protease inhibitor, induces growth arrest, reduced Akt signalling, apoptosis and docetaxel sensitisation in NSCLC cell lines. Br J Cancer 2006;95:1653–62. 10.1038/sj.bjc.6603435.

[17] Micco A Di, Frera G, Lugrin J, Jamilloux Y, Hsu ET, Tardivel A, et al. AIM2 inflammasome is activated by pharmacological disruption of nuclear envelope integrity. Proc Natl Acad Sci U S A 2016;113:E4671–80. 10.1073/pnas.1602419113.

[18] Driessen C, Kraus M, Joerger M, Rosing H, Bader J, Hitz F, et al. Treatment with the HIV protease inhibitor nelfinavir triggers the unfolded protein response and may overcome proteasome inhibitor resistance of multiple myeloma in combination with bortezomib: A phase I trial (SAKK 65/08). Haematologica 2016;101:346–55. 10.3324/haematol.2015.135780.

[19] Brunner TB, Geiger M, Grabenbauer GG, Lang-Welzenbach M, Mantoni TS, Cavallaro A, et al. Phase I trial of the human immunodeficiency virus protease inhibitor nelfinavir and chemoradiation for locally advanced pancreatic cancer. Journal of Clinical Oncology 2008;26:2699–706. 10.1200/JCO.2007.15.2355.

[20] Rengan R, Mick R, Pryma DA, Lin LL, Christodouleas J, Plastaras JP, et al. Clinical Outcomes of the HIV Protease Inhibitor Nelfinavir with Concurrent Chemoradiotherapy for Unresectable Stage IIIA/IIIB Non-Small Cell Lung Cancer: A Phase 1/2 Trial. JAMA Oncol 2019;5:1464–72. 10.1001/jamaoncol.2019.2095.

[21] Alonso-Basanta M, Fang P, Maity A, Hahn SM, Lustig RA, Dorsey JF. A phase i study of nelfinavir concurrent with temozolomide and radiotherapy in patients with glioblastoma multiforme. J Neurooncol 2014;116:365–72. 10.1007/s11060-013-1303-3.

[22] Agrawal I, Saxena S, Nair P, Jha D, Jha S. Obtaining human microglia from adult human brain tissue. Journal of Visualized Experiments 2020;2020. 10.3791/61438.

[23] Sharma N, Saxena S, Agrawal I, Singh S, Srinivasan V, Arvind S, et al. Differential Expression Profile of NLRs and AIM2 in Glioma and Implications for NLRP12 in Glioblastoma. Sci Rep 2019;9. 10.1038/s41598-019-44854-4.

[24] Agrawal I, Sharma N, Saxena S, Arvind S, Chakraborty D, Chakraborty DB, et al. Dopamine induces functional extracellular traps in microglia. IScience 2021;24. 10.1016/j.isci.2020.101968.

[25] Xu Y, Sun Q, Yuan F, Dong H, Zhang H, Geng R, et al. RND2 attenuates apoptosis and autophagy in glioblastoma cells by targeting the p38 MAPK signalling pathway. Journal of Experimental and Clinical Cancer Research 2020;39. 10.1186/s13046-020-01671-2.

[26] Meena D, Shivakumar D, Rajkhowa S, Bhattacharya N, Solanki P, Chhipa S, et al. Innate Immune Receptor NLRX1: Potential Modulator of Glioblastoma Pathophysiology 2024. 10.1101/2024.09.19.613932.

[27] Leung BM, Lesher-Perez SC, Matsuoka T, Moraes C, Takayama S. Media additives to promote spheroid circularity and compactness in hanging drop platform. Biomater Sci 2015;3:336–44. 10.1039/c4bm00319e.

[28] Pinto G, Saenz-De-Santa-Maria I, Chastagner P, Perthame E, Delmas C, Toulas C, et al. Patient-derived glioblastoma stem cells transfer mitochondria through tunneling nanotubes in tumor organoids. Biochemical Journal 2021;478:21–39. 10.1042/BCJ20200710.

[29] Ho WY, Yeap SK, Ho CL, Rahim RA, Alitheen NB. Development of Multicellular Tumor Spheroid (MCTS) Culture from Breast Cancer Cell and a High Throughput Screening Method Using the MTT Assay. PLoS One 2012;7. 10.1371/journal.pone.0044640.

[30] Agrawal I, Sharma N, Saxena S, Arvind S, Chakraborty D, Chakraborty DB, et al. Dopamine induces functional extracellular traps in microglia. IScience 2021;24. 10.1016/j.isci.2020.101968.

[31] Markowitz M, Conant M, Hurley A, Schluger R, Duran M, Peterkin J, et al. A Preliminary Evaluation of Nelfinavir Mesylate, an Inhibitor of Human Immunodeficiency Virus (HIV)-1 Protease, to Treat HIV Infection. n.d.

[32] Soprano M, Sorriento D, Rusciano MR, Maione AS, Limite G, Forestieri P, et al. Oxidative stress mediates the antiproliferative effects of nelfinavir in breast cancer cells. PLoS One 2016;11. 10.1371/journal.pone.0155970.

[33] Pyrko P, Kardosh A, Wang W, Xiong W, Schönthal AH, Chen TC. HIV-1 protease inhibitors nelfinavir and atazanavir induce malignant glioma death by triggering endoplasmic reticulum stress. Cancer Res 2007;67:10920–8. 10.1158/0008-5472.CAN-07-0796.

[34] Jiang W, Mikochik PJ, Ra JH, Lei H, Flaherty KT, Winkler JD, et al. HIV protease inhibitor nelfinavir inhibits growth of human melanoma cells by induction of cell cycle arrest. Cancer Res 2007;67:1221–7. 10.1158/0008-5472.CAN-06-3377.

[35] Lin RZ, Chang HY. Recent advances in three-dimensional multicellular spheroid culture for biomedical research. Biotechnol J 2008;3:1172–84. 10.1002/biot.200700228.

[36] Mueller-Klieser W. invited review Three-dimensional cell cultures: from molecular mechanisms to clinical applications. 1997.

[37] Lin RZ, Chou LF, Chien CCM, Chang HY. Dynamic analysis of hepatoma spheroid formation: Roles of E-cadherin and β1-integrin. Cell Tissue Res 2006;324:411–22. 10.1007/s00441-005-0148-2.

[38] Khaitan D, Chandna S, Arya MB, Dwarakanath BS. Establishment and characterization of multicellular spheroids from a human glioma cell line; implications for tumor therapy. J Transl Med 2006;4. 10.1186/1479-5876-4-12.

[39] De Grandis RA, Santos PW da S dos, Oliveira KM de, Machado ART, Aissa AF, Batista AA, et al. Novel lawsone-containing ruthenium(II) complexes: Synthesis, characterization and anticancer activity on 2D and 3D spheroid models of prostate cancer cells. Bioorg Chem 2019;85:455–68. 10.1016/j.bioorg.2019.02.010.

[40] Tavener AM, Phelps MC, Daniels RL. Anthracycline-induced cytotoxicity in the GL261 glioma model system. Mol Biol Rep 2021;48:1017–23. 10.1007/s11033-020-06109-8.

[41] Amiri M, Basiri M, Eskandary H, Akbarnejad Z, Esmaeeli M, Masoumi-Ardakani Y, et al. Cytotoxicity of carboplatin on human glioblastoma cells is reduced by the concomitant exposure to an extremely low-frequency electromagnetic field (50 Hz, 70 G). Electromagn Biol Med 2018;37:138–45. 10.1080/15368378.2018.1477052.

[42] Kciuk M, Gielecińska A, Mujwar S, Kołat D, Kałuzińska-Kołat Ż, Celik I, et al. Doxorubicin—An Agent with Multiple Mechanisms of Anticancer Activity. Cells 2023;12. 10.3390/cells12040659.

[43] Rabik CA, Dolan ME. Molecular Mechanisms of Resistance and Toxicity Associated with Platinating Agents. n.d.

[44] Sun L, Niu L, Zhu X, Hao J, Wang P, Wang H. Antitumour effects of a protease inhibitor, nelfinavir, in hepatocellular carcinoma cancer cells. Journal of Chemotherapy 2012;24:161–6. 10.1179/1973947812Y.0000000011.

[45] Yang Y, Ikezoe T, Nishioka C, Bandobashi K, Takeuchi T, Adachi Y, et al. NFV, an HIV-1 protease inhibitor, induces growth arrest, reduced Akt signalling, apoptosis and docetaxel sensitisation in NSCLC cell lines. Br J Cancer 2006;95:1653–62. 10.1038/sj.bjc.6603435.

[46] Yang Y, Ikezoe T, Takeuchi T, Adachi Y, Ohtsuki Y, Takeuchi S, et al. HIV-1 protease inhibitor induces growth arrest and apoptosis of human prostate cancer LNCaP cells in vitro and in vivo in conjunction with blockade of androgen receptor STAT3 and AKT signaling. Cancer Sci 2005;96:425–33. 10.1111/j.1349-7006.2005.00063.x.

[47] Crowley LC, Marfell BJ, Scott AP, Waterhouse NJ. Quantitation of apoptosis and necrosis by annexin V binding, propidium iodide uptake, and flow cytometry. Cold Spring Harb Protoc 2016;2016:953–7. 10.1101/pdb.prot087288.

[48] Raucci A, Palumbo R, Bianchi ME. HMGB1: A signal of necrosis. Autoimmunity 2007;40:285–9. 10.1080/08916930701356978.

[49] Sun Q, Loughran P, Shapiro R, Shrivastava IH, Antoine DJ, Li T, et al. Redox-dependent regulation of hepatocyte absent in melanoma 2 inflammasome activation in sterile liver injury in mice. Hepatology 2017;65:253–68. 10.1002/hep.28893.

[50] Patil S, Sengupta K. Role of A- and B-type lamins in nuclear structure–function relationships. Biol Cell 2021;113:295–310. 10.1111/boc.202000160.

[51] Gaidt MM, Ebert TS, Chauhan D, Schmidt T, Schmid-Burgk JL, Rapino F, et al. Human Monocytes Engage an Alternative Inflammasome Pathway. Immunity 2016;44:833–46. 10.1016/j.immuni.2016.01.012.

[52] Chen PA, Shrivastava G, Balcom EF, McKenzie BA, Fernandes J, Branton WG, et al. Absent in melanoma 2 regulates tumor cell proliferation in glioblastoma multiforme. J Neurooncol 2019;144:265–73. 10.1007/s11060-019-03230-y.

[53] Fernandes-Alnemri T, Yu JW, Datta P, Wu J, Alnemri ES. AIM2 activates the inflammasome and cell death in response to cytoplasmic DNA. Nature 2009;458:509–13. 10.1038/nature07710.

[54] Sharma M, de Alba E. Assembly mechanism of the inflammasome sensor AIM2 revealed by single molecule analysis. Nat Commun 2023;14. 10.1038/s41467-023-43691-4.

[55] Gieryng A, Pszczolkowska D, Walentynowicz KA, Rajan WD, Kaminska B. Immune microenvironment of gliomas. Laboratory Investigation 2017;97:498–518. 10.1038/labinvest.2017.19.

[56] Roth P, Junker M, Tritschler I, Mittelbronn M, Dombrowski Y, Breit SN, et al. GDF-15 contributes to proliferation and immune escape of malignant gliomas. Clinical Cancer Research 2010;16:3851–9. 10.1158/1078-0432.CCR-10-0705.

[57] Shabtay-Orbach A, Amit M, Binenbaum Y, Na’Ara S, Gil Z. Paracrine regulation of glioma cells invasion by astrocytes is mediated by glial-derived neurotrophic factor. Int J Cancer 2015;137:1012–20. 10.1002/ijc.29380.

[58] Molina ML, García-Bernal D, Martinez S, Valdor R. Autophagy in the immunosuppressive perivascular microenvironment of glioblastoma. Cancers (Basel) 2020;12. 10.3390/cancers12010102.

[59] Hambardzumyan D, Gutmann DH, Kettenmann H. The role of microglia and macrophages in glioma maintenance and progression. Nat Neurosci 2015;19:20–7. 10.1038/nn.4185.

[60] Ogawa J, Pao GM, Shokhirev MN, Verma IM. Glioblastoma Model Using Human Cerebral Organoids. Cell Rep 2018;23:1220–9. 10.1016/j.celrep.2018.03.105.

[61] Hubert CG, Rivera M, Spangler LC, Wu Q, Mack SC, Prager BC, et al. A three-dimensional organoid culture system derived from human glioblastomas recapitulates the hypoxic gradients and cancer stem cell heterogeneity of tumors found in vivo. Cancer Res 2016;76:2465–77. 10.1158/0008-5472.CAN-15-2402.

[62] Chen L, Zhang Y, Yang J, Hagan JP, Li M. Vertebrate animal models of glioma: Understanding the mechanisms and developing new therapies. Biochim Biophys Acta Rev Cancer 2013;1836:158–65. 10.1016/j.bbcan.2013.04.003.

[63] Zhou Z hua, Ping Y fang, Yu S cang, Yi L, Yao X hong, Chen J hong, et al. A novel approach to the identification and enrichment of cancer stem cells from a cultured human glioma cell line. Cancer Lett 2009;281:92–9. 10.1016/j.canlet.2009.02.033.

[64] Bertotti A, Migliardi G, Galimi F, Sassi F, Torti D, Isella C, et al. A molecularly annotated platform of patient-derived xenografts (“xenopatients”) identifies HER2 as an effective therapeutic target in cetuximab-resistant colorectal cancer. Cancer Discov 2011;1:508–23. 10.1158/2159-8290.CD-11-0109.

[65] Gao H, Korn JM, Ferretti S, Monahan JE, Wang Y, Singh M, et al. High-throughput screening using patient-derived tumor xenografts to predict clinical trial drug response. Nat Med 2015;21:1318–25. 10.1038/nm.3954.

[66] Ben-David U, Ha G, Tseng YY, Greenwald NF, Oh C, Shih J, et al. Patient-derived xenografts undergo mouse-specific tumor evolution. Nat Genet 2017;49:1567–75. 10.1038/ng.3967.

[67] Jacob F, Salinas RD, Zhang DY, Nguyen PTT, Schnoll JG, Wong SZH, et al. A Patient-Derived Glioblastoma Organoid Model and Biobank Recapitulates Inter-and Intra-tumoral Heterogeneity. Cell 2020;180:188–204.e22. 10.1016/j.cell.2019.11.036.

[68] Chen CC, Li HW, Wang YL, Lee CC, Shen YC, Hsieh CY, et al. Patient-derived tumor organoids as a platform of precision treatment for malignant brain tumors. Sci Rep 2022;12. 10.1038/s41598-022-20487-y.

[69] Choubey D, Walter S, Geng Y, Xin H. Cytoplasmic localization of the interferon-inducible protein that is encoded by the AIM2 (absent in melanoma) gene from the 200-gene family. FEBS Lett 2000;474:38–42. 10.1016/S0014-5793(00)01571-4.

[70] Lugrin J, Martinon F. The AIM2 inflammasome: Sensor of pathogens and cellular perturbations. Immunol Rev 2018;281:99–114. 10.1111/imr.12618.

[71] Bürckstümmer T, Baumann C, Blüml S, Dixit E, Dürnberger G, Jahn H, et al. An orthogonal proteomic-genomic screen identifies AIM2 as a cytoplasmic DNA sensor for the inflammasome. Nat Immunol 2009;10:266–72. 10.1038/ni.1702.

[72] Hornung V, Ablasser A, Charrel-Dennis M, Bauernfeind F, Horvath G, Caffrey DR, et al. AIM2 recognizes cytosolic dsDNA and forms a caspase-1-activating inflammasome with ASC. Nature 2009;458:514–8. 10.1038/nature07725.

[73] Albrecht M, Choubey D, Lengauer T. The HIN domain of IFI-200 proteins consists of two OB folds. Biochem Biophys Res Commun 2005;327:679–87. 10.1016/j.bbrc.2004.12.056.

[74] Lu A, Kabaleeswaran V, Fu T, Magupalli VG, Wu H. Crystal structure of the F27G AIM2 PYD mutant and similarities of its self-association to DED/DED interactions. J Mol Biol 2014;426:1420–7. 10.1016/j.jmb.2013.12.029.

[75] Choubey D. Absent in melanoma 2 proteins in the development of cancer. Cellular and Molecular Life Sciences 2016;73:4383–95. 10.1007/s00018-016-2296-9.

[76] Deyoung KL, Ray ME, Su YA, Anzick SL, Johnstone RW, Trapani JA, et al. Cloning a novel member of the human interferon-inducible gene family associated with control of tumorigenicity in a model of human melanoma. 1997.

[77] Chai D, Shan H, Wang G, Li H, Fang L, Song J, et al. AIM2 is a potential therapeutic target in human renal carcinoma and suppresses its invasion and metastasis via enhancing autophagy induction. Exp Cell Res 2018;370:561–70. 10.1016/j.yexcr.2018.07.021.

[78] Ma X, Guo P, Qiu Y, Mu K, Zhu L, Zhao W, et al. Loss of AIM2 expression promotes hepatocarcinoma progression through activation of mTOR-S6K1 pathway. vol. 7. n.d.

[79] Chen LC, Wang LJ, Tsang NM, Ojcius DM, Chen CC, Ouyang CN, et al. Tumour inflammasome-derived IL-1β recruits neutrophils and improves local recurrence-free survival in EBV-induced nasopharyngeal carcinoma. EMBO Mol Med 2012;4:1276–93. 10.1002/emmm.201201569.

[80] Kondo Y, Nagai K, Nakahata S, Saito Y, Ichikawa T, Suekane A, et al. Overexpression of the DNA sensor proteins, absent in melanoma 2 and interferon-inducible 16, contributes to tumorigenesis of oral squamous cell carcinoma with p53 inactivation. Cancer Sci 2012;103:782–90. 10.1111/j.1349-7006.2012.02211.x.

[81] Zhang M, Jin C, Yang Y, Wang K, Zhou Y, Zhou Y, et al. AIM2 promotes non-small-cell lung cancer cell growth through inflammasome-dependent pathway. J Cell Physiol 2019;234:20161–73. 10.1002/jcp.28617.

[82] Wilson JE, Petrucelli AS, Chen L, Koblansky AA, Truax AD, Oyama Y, et al. Inflammasome-independent role of AIM2 in suppressing colon tumorigenesis via DNA-PK and Akt. Nat Med 2015;21:906–13. 10.1038/nm.3908.

[83] Puschmann TB, Zandén C, De Pablo Y, Kirchhoff F, Pekna M, Liu J, et al. Bioactive 3D cell culture system minimizes cellular stress and maintains the in vivo-like morphological complexity of astroglial cells. Glia 2013;61:432–40. 10.1002/glia.22446.

[84] Hawkins BT, Grego S, Sellgren KL. Three-dimensional culture conditions differentially affect astrocyte modulation of brain endothelial barrier function in response to transforming growth factor β1. Brain Res 2015;1608:167–76. 10.1016/j.brainres.2015.02.025.

[85] Brunner TB, Geiger M, Grabenbauer GG, Lang-Welzenbach M, Mantoni TS, Cavallaro A, et al. Phase I trial of the human immunodeficiency virus protease inhibitor nelfinavir and chemoradiation for locally advanced pancreatic cancer. Journal of Clinical Oncology 2008;26:2699–706. 10.1200/JCO.2007.15.2355.

[86] Driessen C, Kraus M, Joerger M, Rosing H, Bader J, Hitz F, et al. Treatment with the HIV protease inhibitor nelfinavir triggers the unfolded protein response and may overcome proteasome inhibitor resistance of multiple myeloma in combination with bortezomib: A phase I trial (SAKK 65/08). Haematologica 2016;101:346–55. 10.3324/haematol.2015.135780.

[87] Garcia-Soto AE, McKenzie ND, Whicker ME, Pearson JM, Jimenez EA, Portelance L, et al. Phase 1 trial of nelfinavir added to standard cisplatin chemotherapy with concurrent pelvic radiation for locally advanced cervical cancer. Cancer 2021;127:2279–93. 10.1002/cncr.33449.

[88] Rengan R, Mick R, Pryma D, Rosen MA, Lin LL, Maity AM, et al. A phase i trial of the HIV protease inhibitor nelfinavir with concurrent chemoradiotherapy for unresectable stage IIIA/IIIB non-small cell lung cancer: A report of toxicities and clinical response. Journal of Thoracic Oncology 2012;7:709–15. 10.1097/JTO.0b013e3182435aa6.

[89] Gunti S, Hoke ATK, Vu KP, London NR. Organoid and spheroid tumor models: Techniques and applications. Cancers (Basel) 2021;13:1–18. 10.3390/cancers13040874.

[90] Xu H, Jiao D, Liu A, Wu K. Tumor organoids: applications in cancer modeling and potentials in precision medicine. J Hematol Oncol 2022;15. 10.1186/s13045-022-01278-4.

[91] Verduin M, Hoeben A, De Ruysscher D, Vooijs M. Patient-Derived Cancer Organoids as Predictors of Treatment Response. Front Oncol 2021;11. 10.3389/fonc.2021.641980.

[92] Jones DTW, Kocialkowski S, Liu L, Pearson DM, Bäcklund LM, Ichimura K, et al. Tandem duplication producing a novel oncogenic BRAF fusion gene defines the majority of pilocytic astrocytomas. Cancer Res 2008;68:8673–7. 10.1158/0008-5472.CAN-08-2097.

[93] Yan H, Parsons DW, Jin G, McLendon R, Rasheed BA, Yuan W, et al. IDH1 and IDH2 Mutations in Gliomas. New England Journal of Medicine 2009;360:765–73. 10.1056/nejmoa0808710.

[94] Comprehensive, Integrative Genomic Analysis of Diffuse Lower-Grade Gliomas. New England Journal of Medicine 2015;372:2481–98. 10.1056/NEJMoa1402121.

