## supplementray figures with legend for "Repurposing Nelfinavir: AIM2 Inflammasome-Driven Anti-tumor Effects in Glioblastoma"

#### Figure S1. Cell viability assay in Nelfinavir (4 μM) treated Glioblastoma and astrocyte spheroids, related to Figure 1

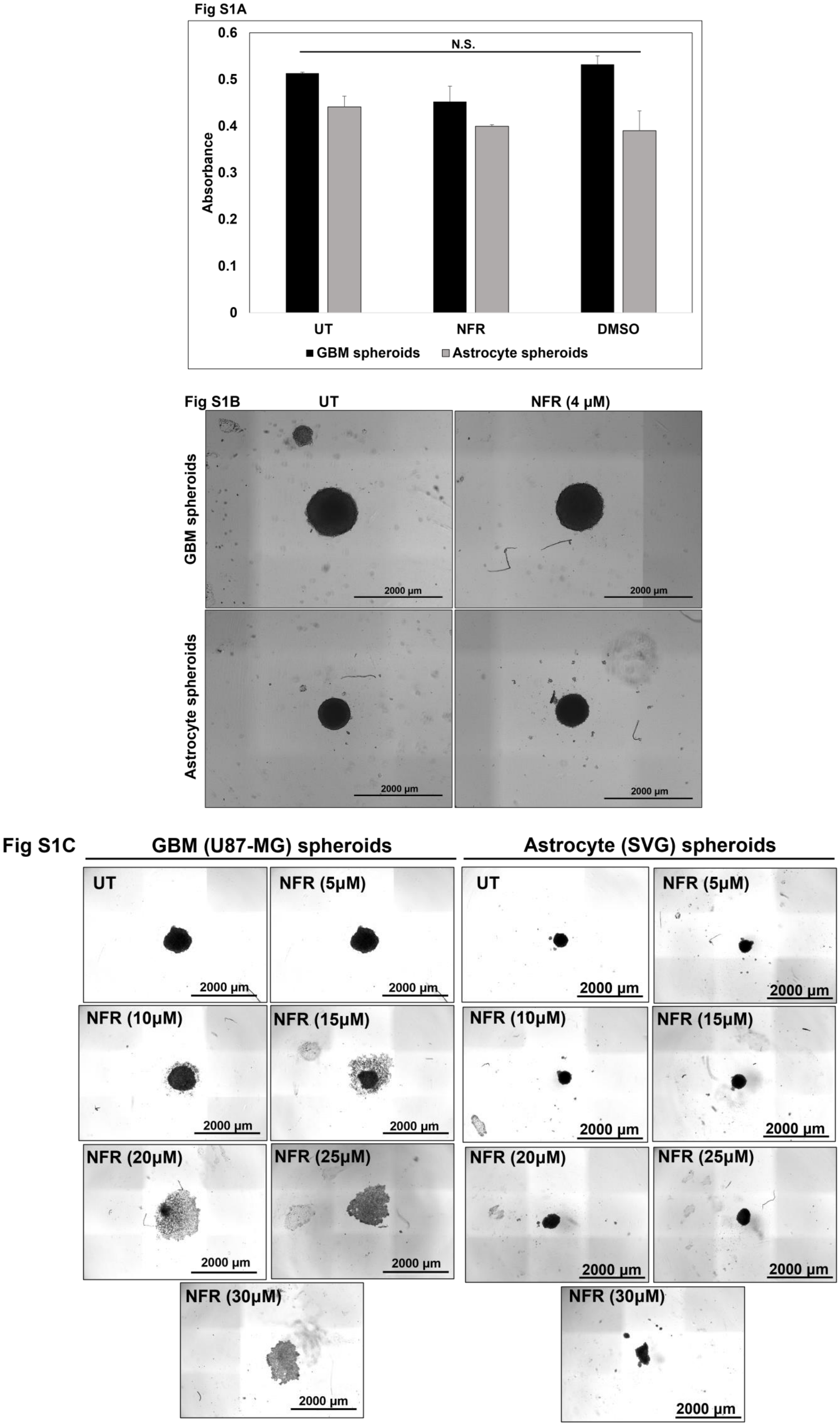

Fig S1D

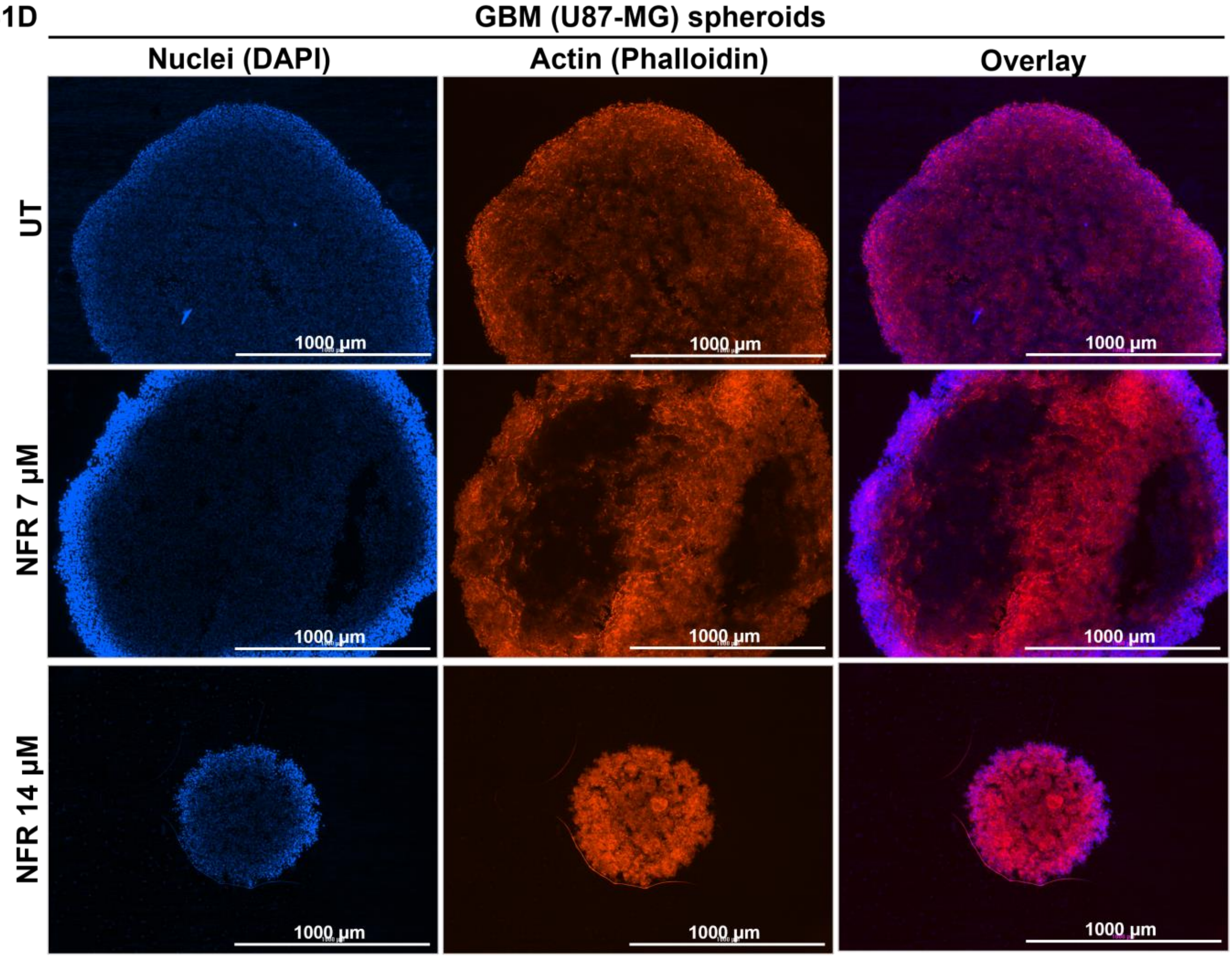


Fig S1E

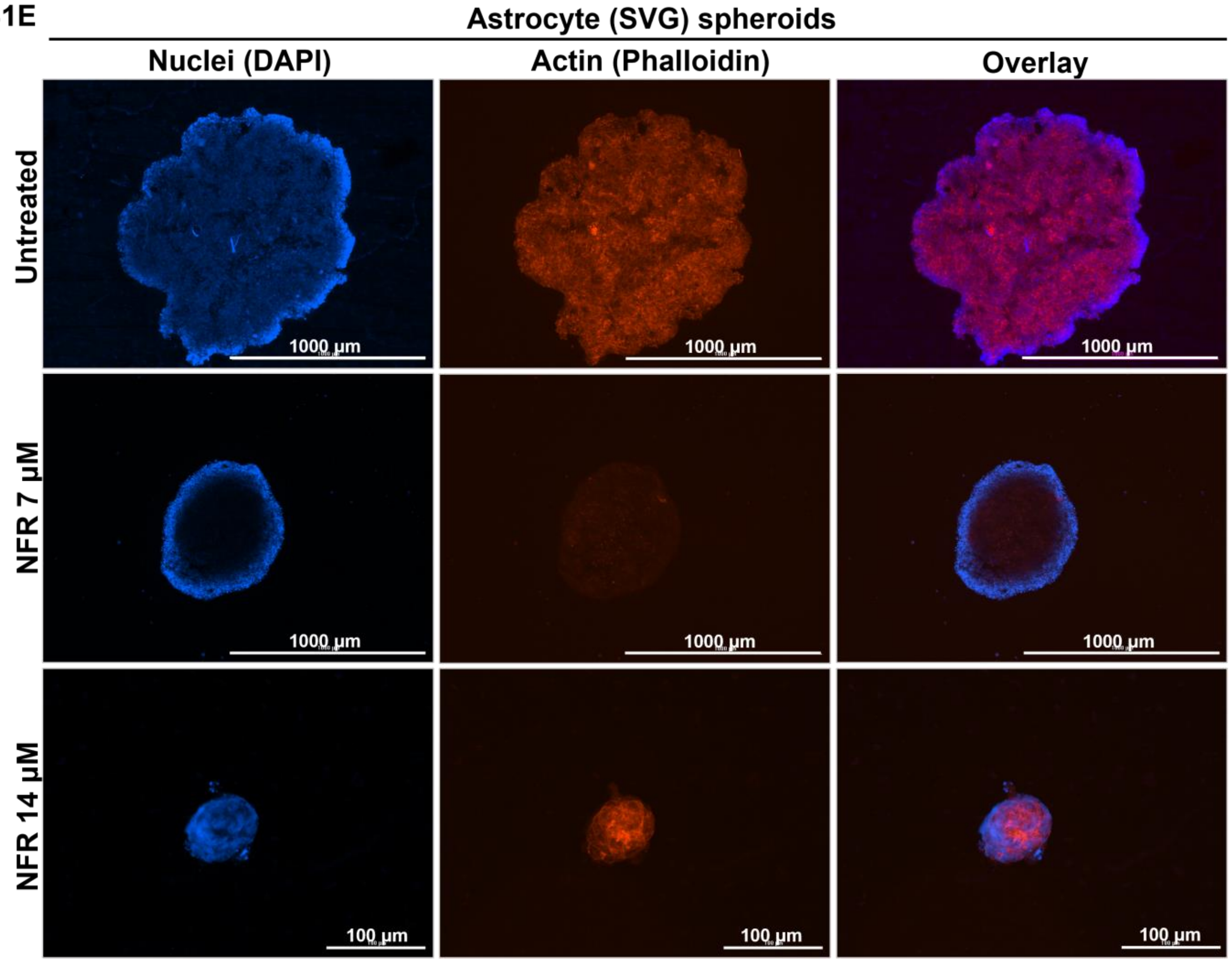


**Figure S1. Cell viability assay in Nelfinavir (4 µM) treated Glioblastoma and astrocyte spheroids, related to Figure 1:** S1A. Cell viability assay was performed in U87-MG and SVG spheroids treated with NFR (4µM) for 48 hours. Data are represented as mean ± SEM. Error bars indicated SEM. N.S. = non-significant. S1B. Brightfield images of U87-MG and SVG spheroids treated with NFR (4µM) for 48 hours. Images are representative of 2 experiments. Scale bar, 2000µm. S1C. Brightfield images of U87-MG and SVG spheroids treated with NFR at indicated concentrations for 48 hours. Images are representative of 2 experiments. Scale bar, 2000µm. S1D-E. UT and NFR (7µM, 14µM, 48 hours) treated U87-MG and SVG spheroids were stained with Phalloidin (actin, red) and DAPI (nuclei, blue). At least 3 frames were imaged per spheroid. Images are representative of all frames. Scale bar,1000 µm.

### Figure S2. Cell viability assay in Nelfinavir (4 µM), CARBO, and DOX-treated Glioblastoma spheroids, related to Figure 2

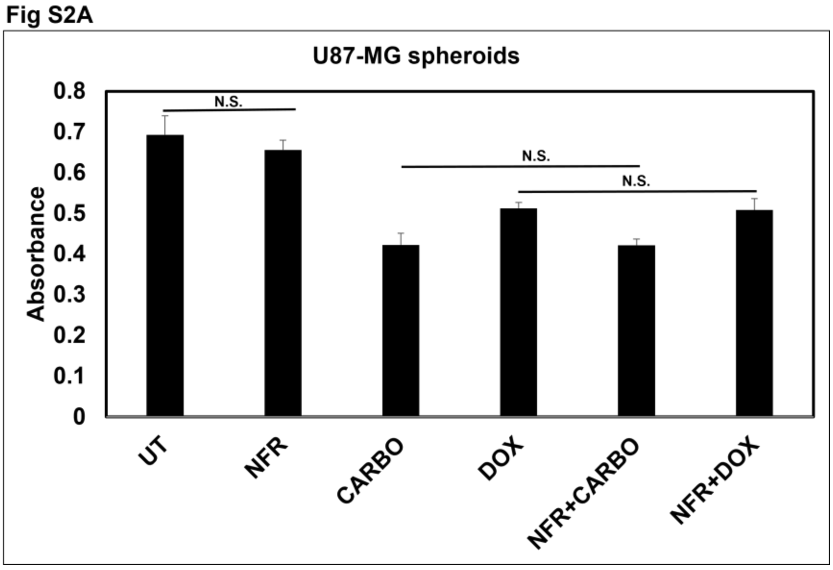


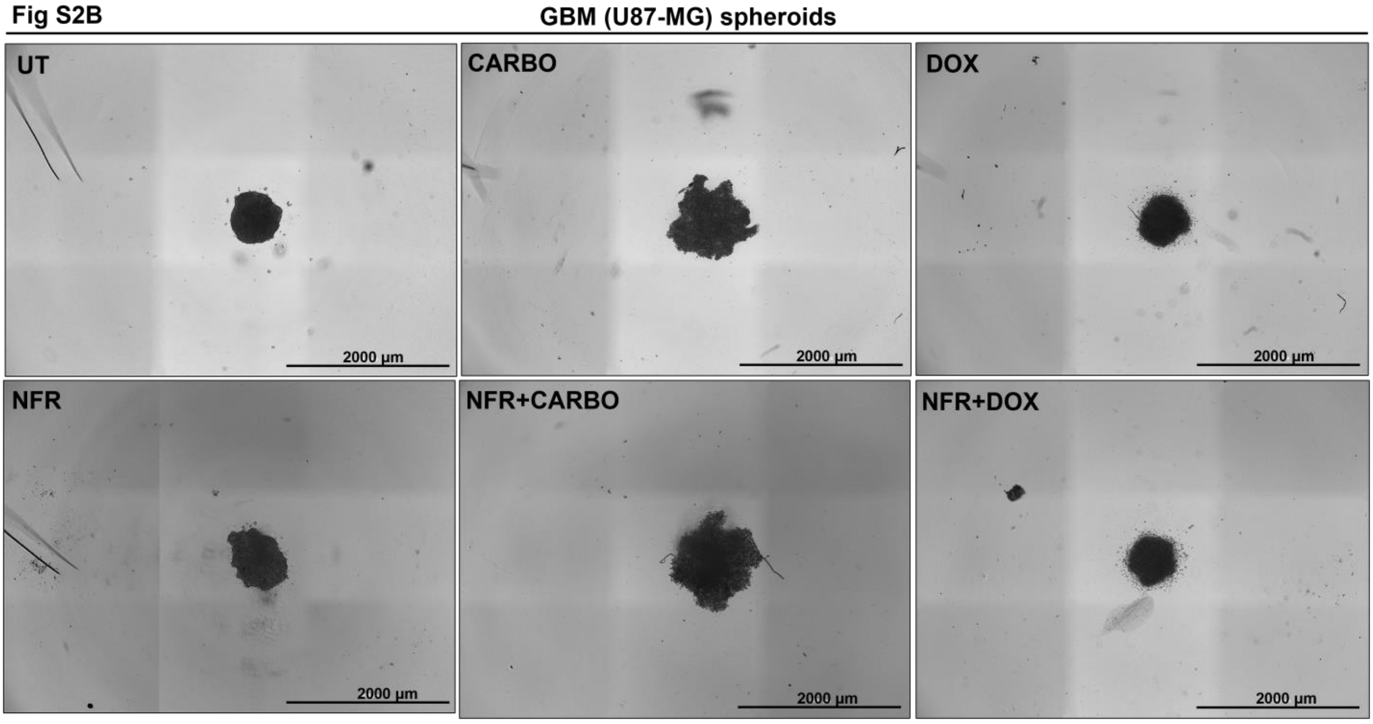


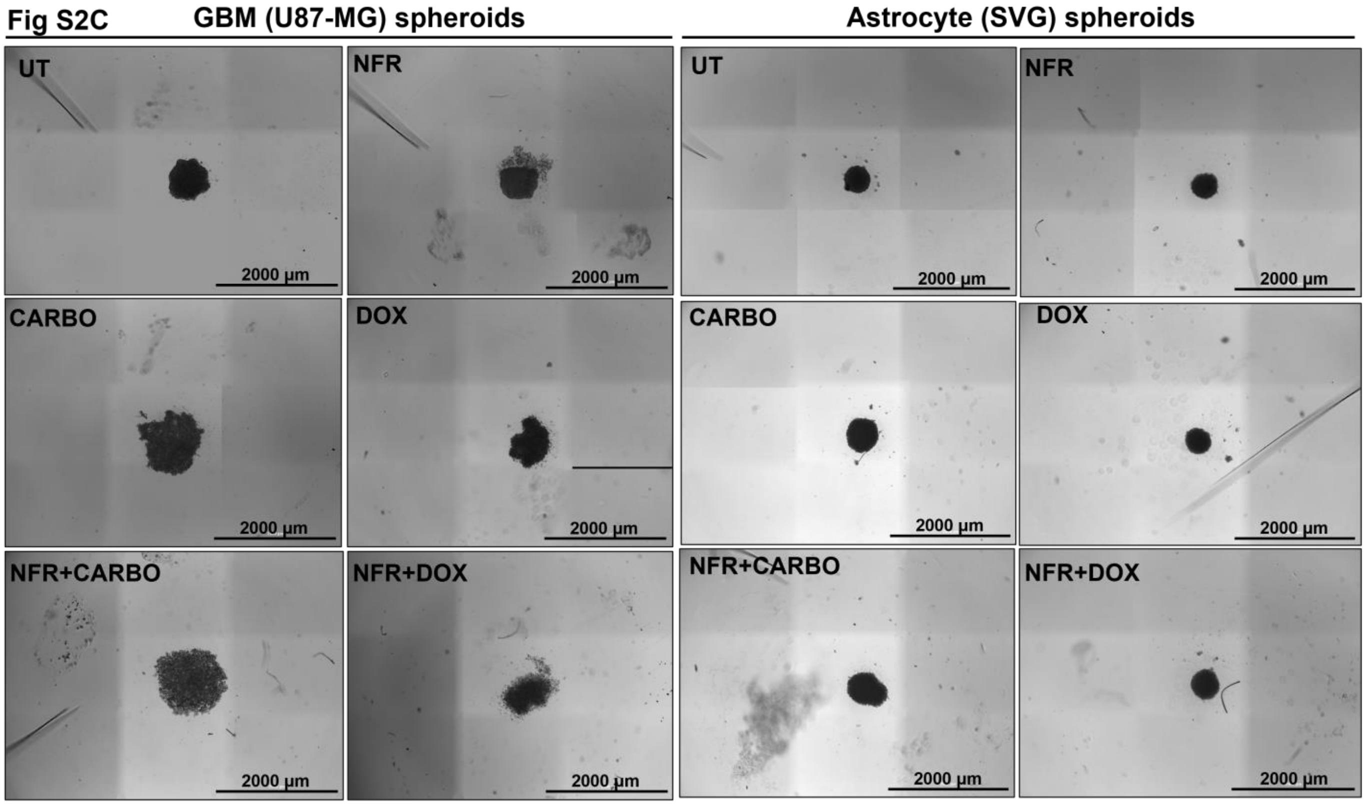


**Figure S2. Cell viability assay in Nelfinavir (4 µM), Carboplatin (CARBO), and Doxorubicin (DOX)-treated Glioblastoma spheroids, related to Figure 2:** S2A. Cell viability assay was performed in U87-MG spheroids treated with NFR (4µM), CARBO (2.7mM), DOX (4.9µM), or in indicated combination for 48 hours. Data are represented as mean ± SEM. Error bars indicated SEM. N.S. = non-significant. S2B. Brightfield images of U87-MG spheroids treated with NFR (4µM), CARBO, DOX, or in indicated combination for 48 hours. Scale bar, 2000µm. S2C. Brightfield images of U87-MG and SVG spheroids treated with NFR (7µM), CARBO, DOX, or in indicated combination for 48 hours. Images are representative of 3 experiments. Scale bar, 2000µm.

### Figure S3. Nelfinavir induces Apoptosis and Necrosis in Glioblastoma cells, related to Figure 3, 4

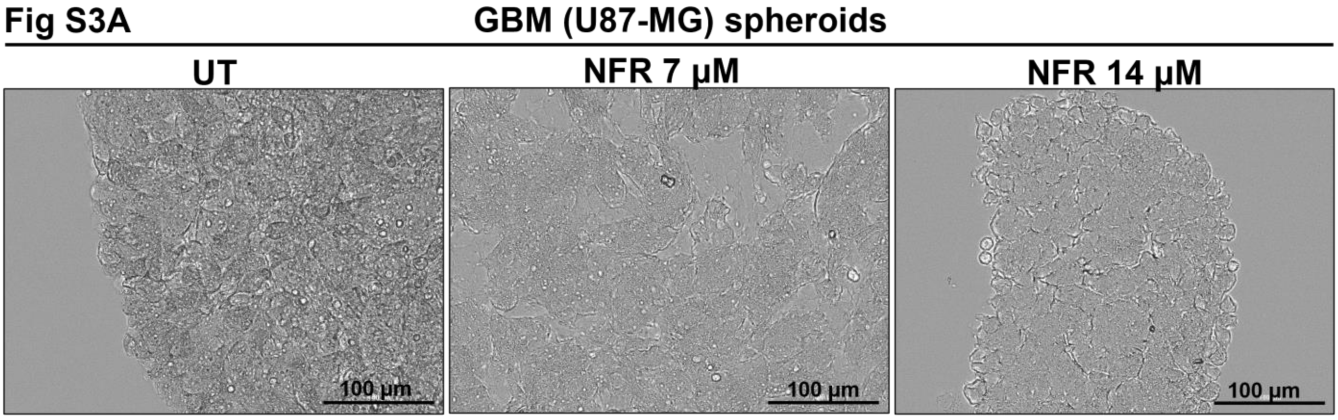


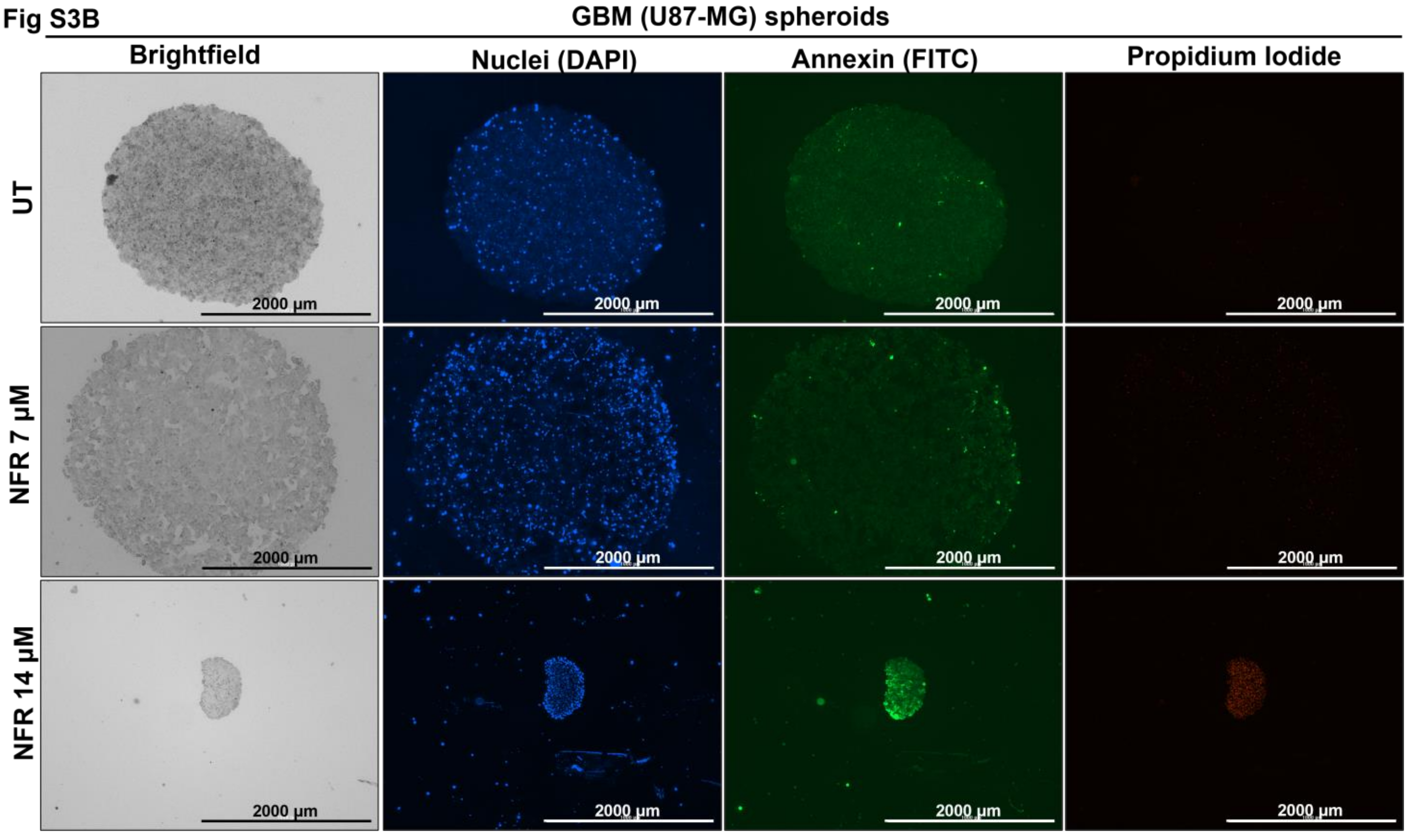


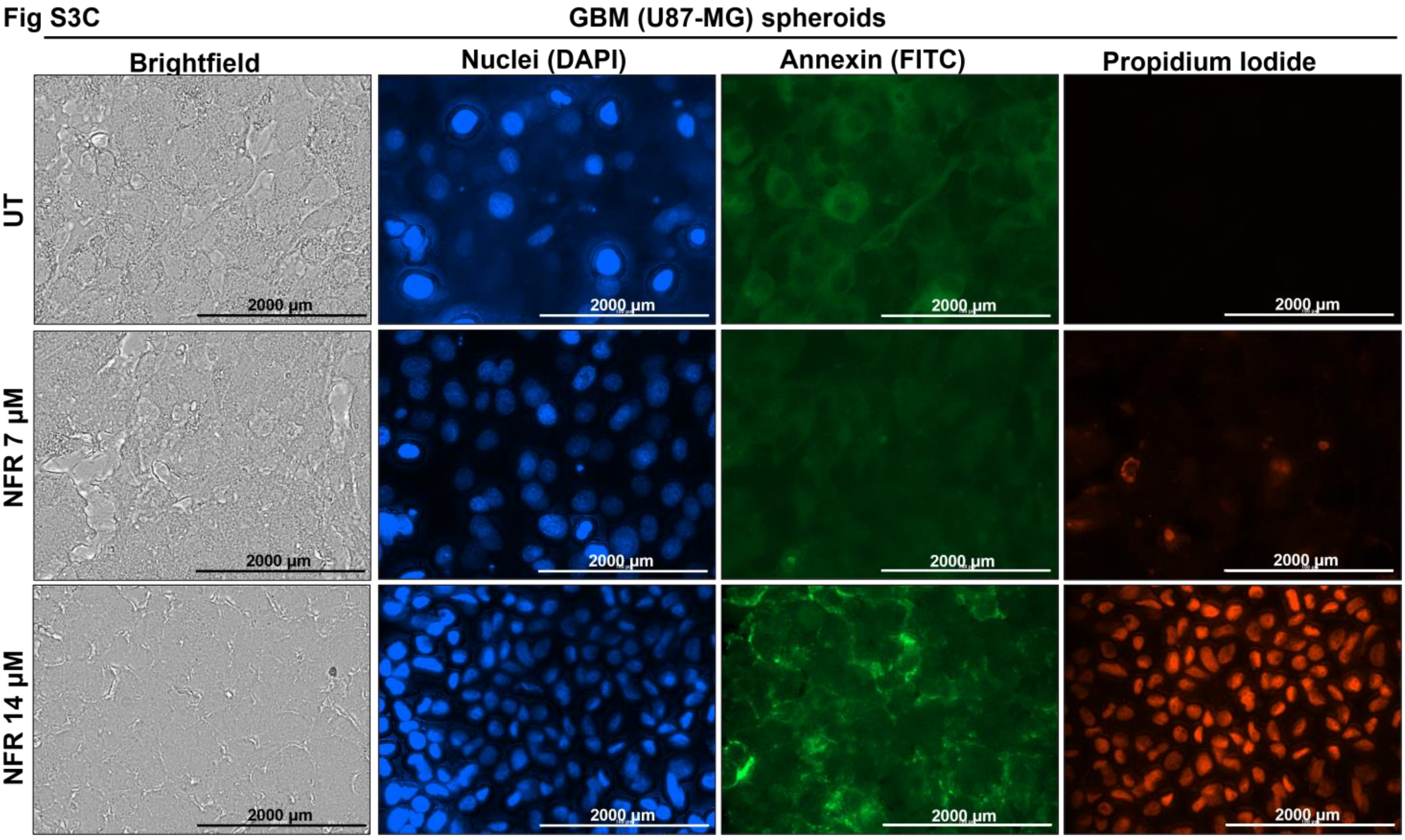

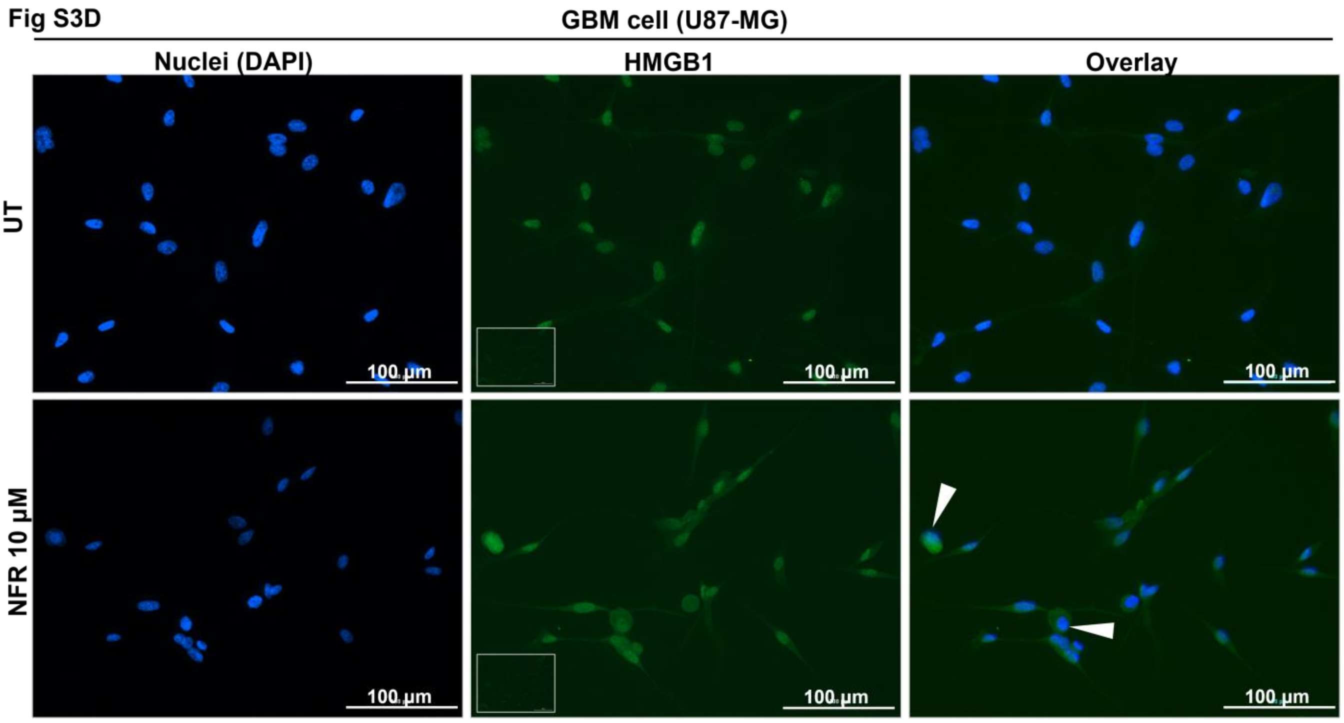


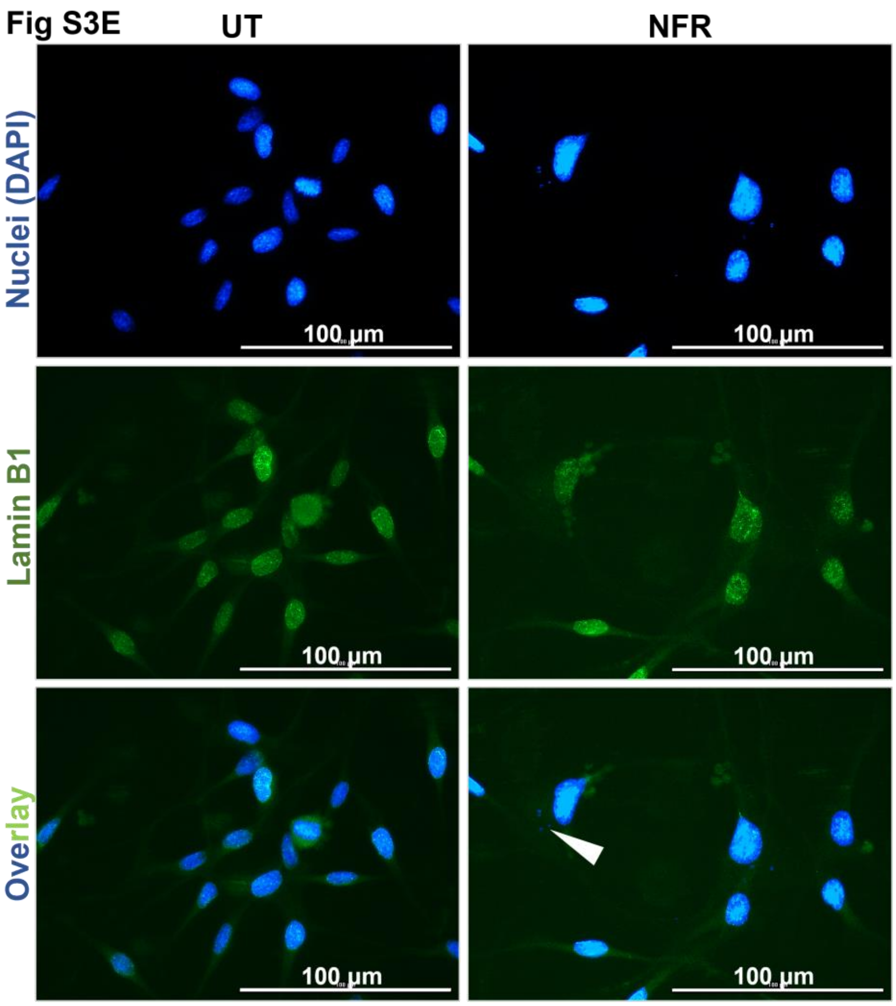


**Figure S3. Nelfinavir induces Apoptosis and Necrosis in Glioblastoma cells, related to Figure 3:** S3A. Brightfield images of UT and NFR (7μM, 14μM, 48 hours) treated U87-MG spheroids. At least 3 frames were imaged per spheroid. Images are representative of all frames. Images were taken at 20X magnification. Scale bar,100 μm. S3B. 4X magnified images of UT and NFR (7μM, 14μM, 48 hours) treated U87-MG spheroids stained with annexin V (green), propidium iodide (PI) (red), and DAPI (nuclei, blue). Scale bar, 2000 μm. S3C. 40X magnified images of UT and NFR (7μM, 14μM, 48 hours) treated U87-MG spheroids stained with annexin V (green), propidium iodide (PI) (red), and DAPI (nuclei, blue). At least 3 frames were imaged per spheroid. Images are representative of all frames. Scale bar, 2000 μm. S4D. UT and NFR (10μM, 24 hours) treated U87-MG cells were stained with anti-HMGB1 antibody (green) and DAPI (nuclei, blue). At least 10 frames were imaged per well of the two-well chamber slides. Images are representative of all frames. Inset represents primary antibody control. Images were taken at 20X magnification. Scale bar,100 μm. S3E. UT and NFR (10μM, 24 hours) treated U87-MG cells were stained with anti-Lamin B1 antibody (green) and DAPI (nuclei, blue). At least 10 frames were imaged per well of the two-well chamber slides. Images are representative of all frames. Inset represents primary antibody control. Images were taken at 40X magnification. Scale bar,100 μm

### Figure S4. Differential expression of AIM2 in Glioblastoma cells, related to figure 5

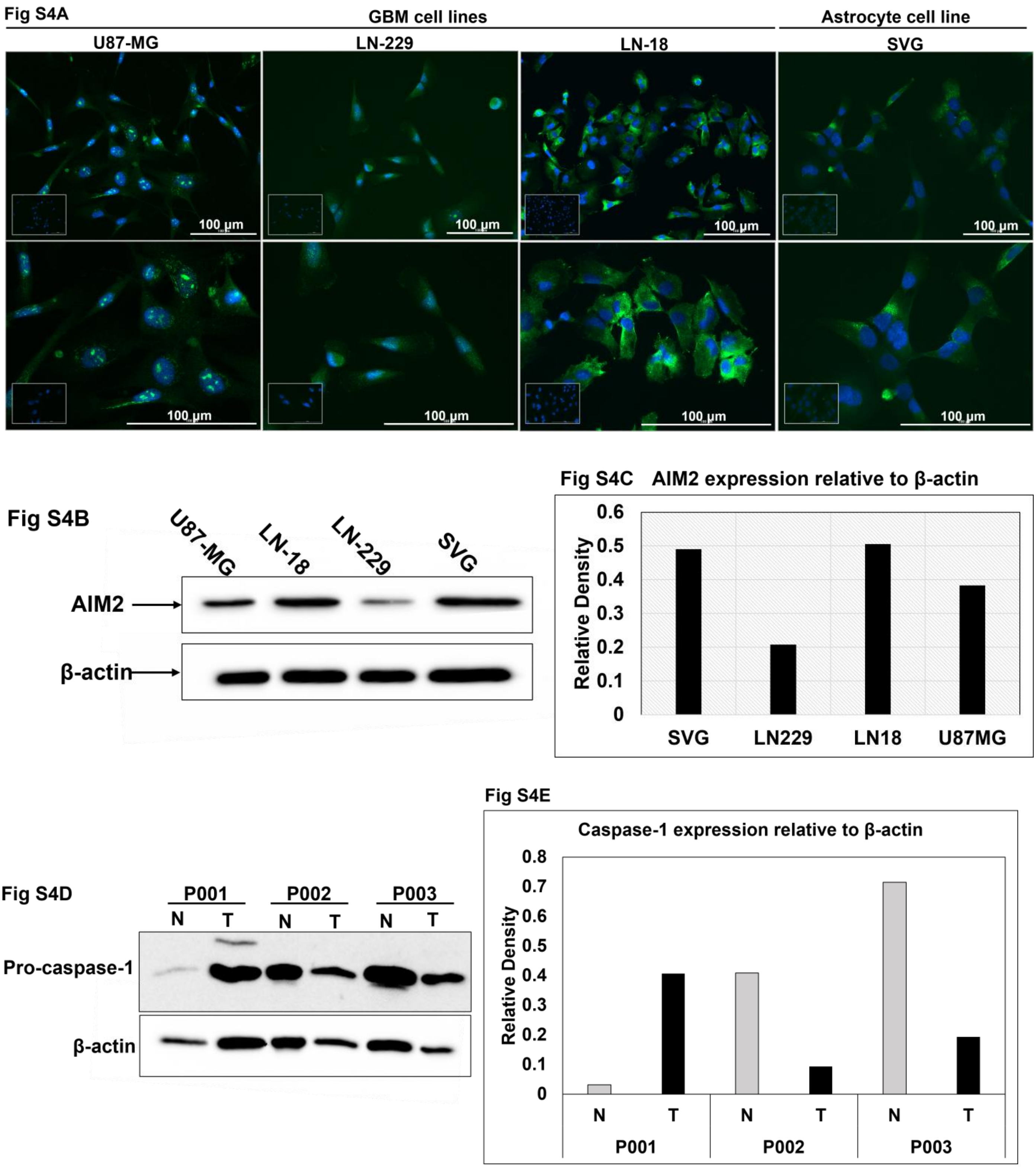


**Figure S4. Differential expression of AIM2 in Glioblastoma cells:** S4A. To check the AIM2 expression across various GBM and astrocyte cell lines, cells were stained with anti-AIM2 antibody (green) and DAPI (nuclei, blue). At least 7 frames were imaged per well of the two-well chamber slides. Images are representative of all frames. Inset represents primary antibody control. Scale bar,100 µm. S4B. AIM2 expression is quantified across various cell types using western blot. β-actin is used as a loading control. 20µg protein sample is loaded in each well. Images are representative of 3 experiments. S4C. The western blot data was quantified using densitometry. The graph is representative of one experiment. S4D. Caspase-1 expression was quantified across diverse glioma-grade patient tissues using western blot. β-actin is used as a loading control (N = normal, T = tumor) (n=3). S4E. The western blot data were quantified using densitometry.

.
